# DNA methylation and survival differences associated with the type of IDH mutation in 1p/19q non-codeleted astrocytomas

**DOI:** 10.1101/2020.12.10.419333

**Authors:** C Mircea S Tesileanu, Wies R Vallentgoed, Marc Sanson, Walter Taal, Paul M Clement, Wolfgang Wick, Alba Ariela Brandes, Jean Francais Baurain, Olivier L Chinot, Helen Wheeler, Sanjeev Gill, Matthew Griffin, Leland Rogers, Roberta Rudà, Michael Weller, Catherine McBain, Jaap Reijneveld, Roelien H Enting, Francesca Caparrotti, Thierry Lesimple, Susan Clenton, Anja Gijtenbeek, Elisabeth Lim, Filip de Vos, Paul J Mulholland, Martin J B Taphoorn, Iris de Heer, Youri Hoogstrate, Maurice de Wit, Lorenzo Boggiani, Sanne Venneker, Jan Oosting, Judith VMG Bovée, Sara Erridge, Michael A Vogelbaum, Anna K Nowak, Warren P Mason, Johan M Kros, Pieter Wesseling, Ken Aldape, Robert B Jenkins, Hendrikus J Dubbink, Brigitta Baumert, Vassilis Golfinopoulos, Thierry Gorlia, Martin van den Bent, Pim J French

## Abstract

Somatic mutations in the isocitrate dehydrogenase genes *IDH1* and *IDH2* occur at high frequency in several tumour types. Even though these mutations are confined to distinct hotspots, we show that gliomas are the only tumour type with an exceptionally high percentage of IDH1^R132H^ mutations. This high prevalence is important as IDH1^R132H^ is presumed to be relatively poor at producing D-2-hydroxyglutarate (D-2HG) whereas high concentrations of this oncometabolite are required to inhibit TET2 DNA demethylating enzymes. Indeed, patients harbouring IDH1^R132H^ mutated tumours have lower levels of genome-wide DNA-methylation, and an associated increased gene expression, compared to tumours with other IDH1/2 mutations (“non-R132H mutations”). This reduced methylation is seen in multiple tumour types and thus appears independent of site of origin. For 1p/19q non-codeleted glioma patients, we show that this difference is clinically relevant: in samples of the randomised phase III CATNON trial, patients harbouring non-R132H mutated tumours have better outcome (HR 0.41, 95% CI [0.24, 0.71], p=0.0013). Non-R132H mutated tumours also had a significantly lower proportion of tumours assigned to prognostically poor DNA-methylation classes (p<0.001). IDH mutation-type was independent in a multivariable model containing known clinical and molecular prognostic factors. To confirm these observations, we validated the prognostic effect of IDH mutation type on a large independent dataset. The observation that non-R132H mutated 1p/19q non-codeleted gliomas have a more favourable prognosis than their IDH1^R132H^ mutated counterpart is clinically relevant and should be taken into account for patient prognostication.

**Single sentence summary:** Astrocytoma patients with tumours harbouring IDH mutations other than p.R132H have increased DNA methylation levels and longer survival

## Introduction

Somatic mutations in the isocitrate dehydrogenase genes *IDH1* and *IDH2* occur at high frequency in various tumour types including gliomas (primary malignant central nervous system tumours), intrahepatic cholangiocarcinomas (bile duct tumours), enchondromas and chondrosarcomas (bone tumours), sinonasal undifferentiated carcinomas and leukemias^1,2^. More sporadic but similar mutations have been found in a wide variety of other tumour types including melanoma, prostate and pancreatic cancer^3^. *IDH1/2* mutations are causal for the disease and tumours often remain dependent on the mutation for growth^4,5^. The importance of the mutation is confirmed by the activity of IDH-inhibitors: inhibiting the mutant activity of either *IDH1* or *IDH2* shows anti-tumour activity in relapsed/refractory *IDH1/2* mutated acute myeloid leukemia^6,7^ and cholangiocarcinoma patients^8^. The objective response rates in these trials are in the order of 40%, though patients eventually relapse. In gliomas however, mutant *IDH1/2* inhibitors have thus far not shown a survival benefit, but further studies on early-stage tumours are ongoing ^9^.

The *IDH1/2* mutations are confined to defined hotspots within the genes that affect either arginine 132 (R132) in *IDH1* or the arginines R172 or R140 in *IDH2*. These mutations change the activity of the wild-type (wt) protein from an enzyme that produces alpha-ketoglutarate (aKG) to an enzyme that produces D-2 hydroxyglutarate (D-2HG)^1,10^. D-2HG in its turn is a main effector in oncogenesis e.g. by inhibiting aKG-dependent dioxygenases, which keeps cells in an undifferentiated state^11,12^. Although *IDH1/2* mutations are confined to these three hotspots, several reports have shown that the IDH-mutation spectrum differs per tumour type^1,13–15^. This difference is interesting as other groups have shown that mutations differ in their ability to produce D-2HG^16,17^. IDH1^R132H^, the *IDH1/2* mutation with relatively low D-2HG production capacity, is the most common mutation in gliomas; other mutations such as IDH1^R132C^ have 10-fold lower *K*_M_ and have higher enzymatic efficiency^16,17^. The differential D-2HG production capacity is supported by observations from cell lines and clinical samples where tumours harbouring the IDH1^R132H^ mutation have lower D-2HG levels compared to those with other IDH mutations^16,18,19^. This difference may have biological implications as not all aKG-dependent enzymes are equally well inhibited by D-2HG ^20,21^.

Here, we have used data from six large and independent DNA methylation datasets (the randomised phase III CATNON clinical trial on anaplastic 1p/19q non-codeleted gliomas^22^, the TCGA-LGG cohort^23^, samples included in the TAVAREC randomised phase 2 clinical trial on 1p/19q non-codeleted gliomas^24^, a large cohort of acute myeloid leukemias (AML)^25^ and a cohort of chondrosarcomas (Venneker et al, accepted for publication)) derived from four different tumour types, to examine the molecular effects of different types of *IDH1/2* mutations. We report that tumours harbouring IDH1^R132H^ mutations, regardless of tumour type, have lower genome-wide DNA methylation levels compared to those harbouring other (‘non-R132H’) *IDH1/2* hotspot mutations. For 1p/19q non-codeleted glioma patients, we show this difference has clinical relevance as patients harbouring such non-R132H mutated tumours have improved survival. Our data support the notion that increased genome-wide DNA methylation levels are associated with improved outcome in this tumour type and indicate that the type of *IDH1/2* mutation should be taken into account for prognostication of 1p/19q non-codeleted glioma patients.

## Methods

### Datasets

The COSMIC database (assessed 27 December 2019) was screened for hotspot *IDH1* (R132) and *IDH2* (R172 and R140) mutations. Mutations were stratified by tumour type; tumours with low prevalence of mutations were concatenated (‘other tumours’: prostate n=11, pancreas n=6, skin n=32, large intestine n=1, soft tissue n=22, endometrium n=1, breast n=9, urinary tract n=2, liver n=7, stomach n=1, upper aerodigestive tract n=35, salivary gland n=1, thyroid n=1). CATNON clinical data^22^ and *IDH1/2* mutation and DNA methylation data (Tesileanu, submitted) were reported previously. TCGA glioma data (DNA methylation and RNA-seq)^23^, MSK-IMPACT data^26^ and AML data^25^ were downloaded from the TCGA data portal. Clinical data and mutation status for the chondrosarcoma data were reported previously (Venneker et al, accepted for publication). Clinical data from the TAVAREC trial were derived from ref ^24^, and supplemented with DNA methylation data of 89 tumours. Most (80%) TAVAREC samples were derived from the initial tumour. Processing of CATNON and TAVAREC DNA methylation data was performed as described (Tesileanu, submitted). For the CATNON, TCGA-astrocytoma and TAVAREC datasets, we included only *IDH1/2* mutated samples from non 1p/19q-codeleted tumours. For *IDH1/2* mutated MSK-IMPACT samples, the distinction between astrocytic and oligodendrocytic tumours was made by absence or presence of telomerase reverse transcriptase (TERT) promoter mutations^27,28^. In the Chinese Glioma Genome Atlas [CGGA] ^29^, the exact IDH-mutation was not noted and therefore limited for the scope of this analysis. We used only the 1p/19q codeleted tumours in this dataset with *IDH2* mutations being designated as “non-R132H” mutations and all *IDH1* mutations as “R132H”. In oligodendrogliomas, IDH1 mutations virtually always result in R132H^14^. RNA-seq data (raw read counts) were normalized and processed using DEseq2.

### Statistical analysis

Survival curves were created using the Kaplan-Meier method. The log-rank test was used to determine survival differences. A Wilcoxon rank test on beta values (i.e. the intensity of the methylated probe/sum of methylated and unmethylated probe intensity) was used to identify differentially methylated probes in CATNON and TCGA-astrocytoma datasets. To increase power in the smaller sized datasets, we performed an *F*-test on M-values (i.e. the log2 ratio of the methylated/unmethylated probe intensities) to identify differentially methylated CpGs using the dmpFinder function in the Minfi Bioconductor package^30^. To further increase statistical power in the chondrosarcoma dataset (required as this dataset had few samples), we first made a selection of the most variable probes (i.e. those with a standard deviation >2; ~5% of the total number of probes) followed by an *F*-test on the M-values. In all differential methylation analysis, p-values were corrected for false discovery rate (adjusted P-value).

Differences in mutation frequencies were determined using a chi squared test. Pathway analysis was performed using Ingenuity pathway analysis (Qiagen, Venlo, the Netherlands). An association model was made with the Cox proportional hazards method and included, next to *IDH1/2* mutation type, factors that are known to be related to outcome from literature such as sex, treatment with temozolomide, age at randomization, WHO performance score, *MGMT* promoter methylation status, use of corticosteroids at randomization, and DNA methylation profiling. All p values below 0.05 were considered significant. Statistical analysis was performed using R version 3.6.3 and packages minfi, stats, rms, survival.

## Results

### The IDH1^R132H^ mutation predominates in gliomas

We screened the catalogue of somatic mutations in cancer (COSMIC) database^31^, extracted *IDH1/2* hotspot mutation data (IDH1^R132^, IDH2^R172^ and IDH2^R140^) and stratified them by tumour organ site. As expected, tumours with a high frequency of *IDH1/2* mutations include central nervous system (CNS), biliary tract, bone, haematopoietic and lymphoid tumours (leukemias). Interestingly, even if there are only three mutational hotspots, there are marked differences in the distribution of mutations between tumour sites (figure 1). For example, the IDH1^R132H^ mutation is by far the most predominant IDH mutation in CNS tumours (n=7265/8026, 90.5%) whereas this mutation is present at much lower frequencies in bone (n=49/361, 13.6%), leukemic (n=519/2995, 17.3%) and other tumours (n=14/129, 10.9%), and thus far has never been identified in biliary tract tumours (n=212) (p<0.001, chi square test). In contrast, the mutation that results in IDH1^R132C^ is quite rare in gliomas (223/8026, 2.8%) but much more prevalent in all other tumour types: bone (n=212/361, 67.1%), leukemic (n=493/2995, 16.5%), biliary tract (n=114/212, 53.8%) and other tumours (n=14/129, 10.9%). This difference is despite the fact that the IDH1^R132H^ and the IDH1^R132C^ are both the result of a transition mutation (G>A and C>T, respectively). In general, transition mutations are much more common than transversion mutations^32^. There is also a major difference in the distribution of *IDH2* mutations which are very common in haematopoietic and lymphoid tumours but rare in all other tumour types. Mutations of the R140 in *IDH2* are virtually exclusive to haematopoietic and lymphoid tumours.

**Figure 1:**
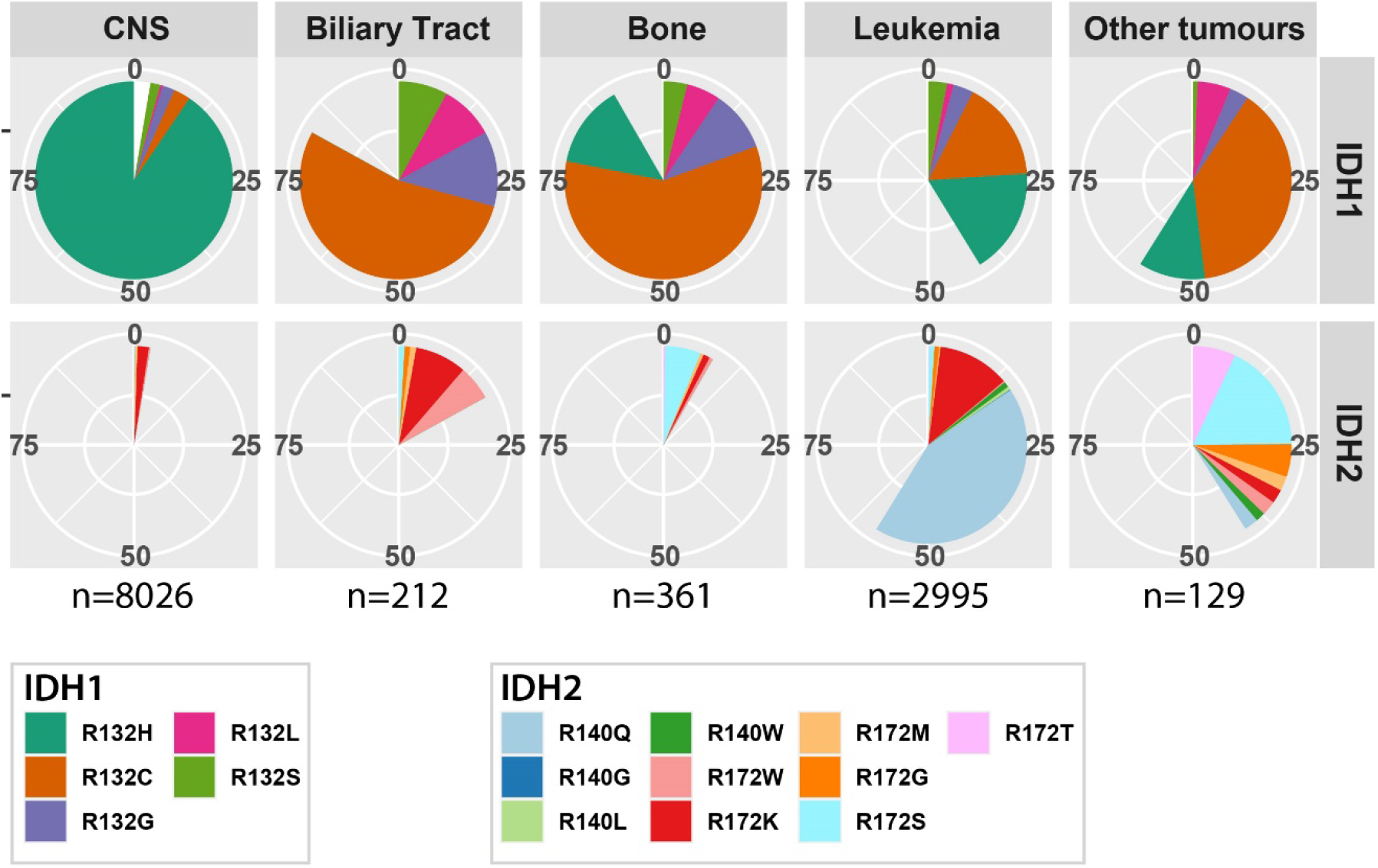
*IDH1* and *IDH2* hotspot mutation distribution separated by site of origin. IDH1^R132H^ mutations are the most predominant mutation in gliomas, IDH2 mutations are most common to haematopoietic tumours.

### DNA methylation is lower in IDH1^R132H^ mutant glioma

Previous reports have shown that D-2HG is a weak inhibitor of TET2 enzymes as relatively high levels of D-2HG are required to inhibit the enzyme^21,33^. We therefore hypothesized that IDH mutations that are presumed to be poor in producing D-2HG (i.e. IDH1^R132H^), produce levels of the oncometabolite that are insufficient to completely inhibit the aKG-dependent dioxygenase TET2. If so, based on the molecular function of TET2 enzymes in mediating the first step in DNA demethylation, IDH1^R132H^ mutated tumours may have lower levels of DNA methylation than those harbouring other hotspot IDH mutations (“non-R132H” mutations).

To test this hypothesis, we used genome-wide DNA methylation data from CATNON trial samples and compared profiles of IDH1^R132H^ mutated tumours (n=369, presumed low D-2HG production) to those harbouring other “non-R132H” *IDH1* and *IDH2* hotspot mutations (n=69, presumed high D-2HG production). Our data shows that the overall level of DNA methylation was significantly lower in tumours harbouring IDH1^R132H^ mutations compared to tumours harbouring non-R132H mutations. For example, there are 2461 probes showing a reduction in beta values > 0.2 in IDH1^R132H^ mutated tumours (at p<0.01) but there are no probes showing an increase > 0.2. This is exemplified in the volcano plot where a strong skew towards increased DNA methylation in non-R132H mutated samples is observed (figure 2A). Probes showing the largest increase in DNA methylation were those that were partially methylated in IDH1^R132H^ mutated tumours (i.e. probes with beta values between 0.25 and 0.75); there were few probes that became (partially) methylated from an unmethylated state (figure 2B).

**Figure 2:**
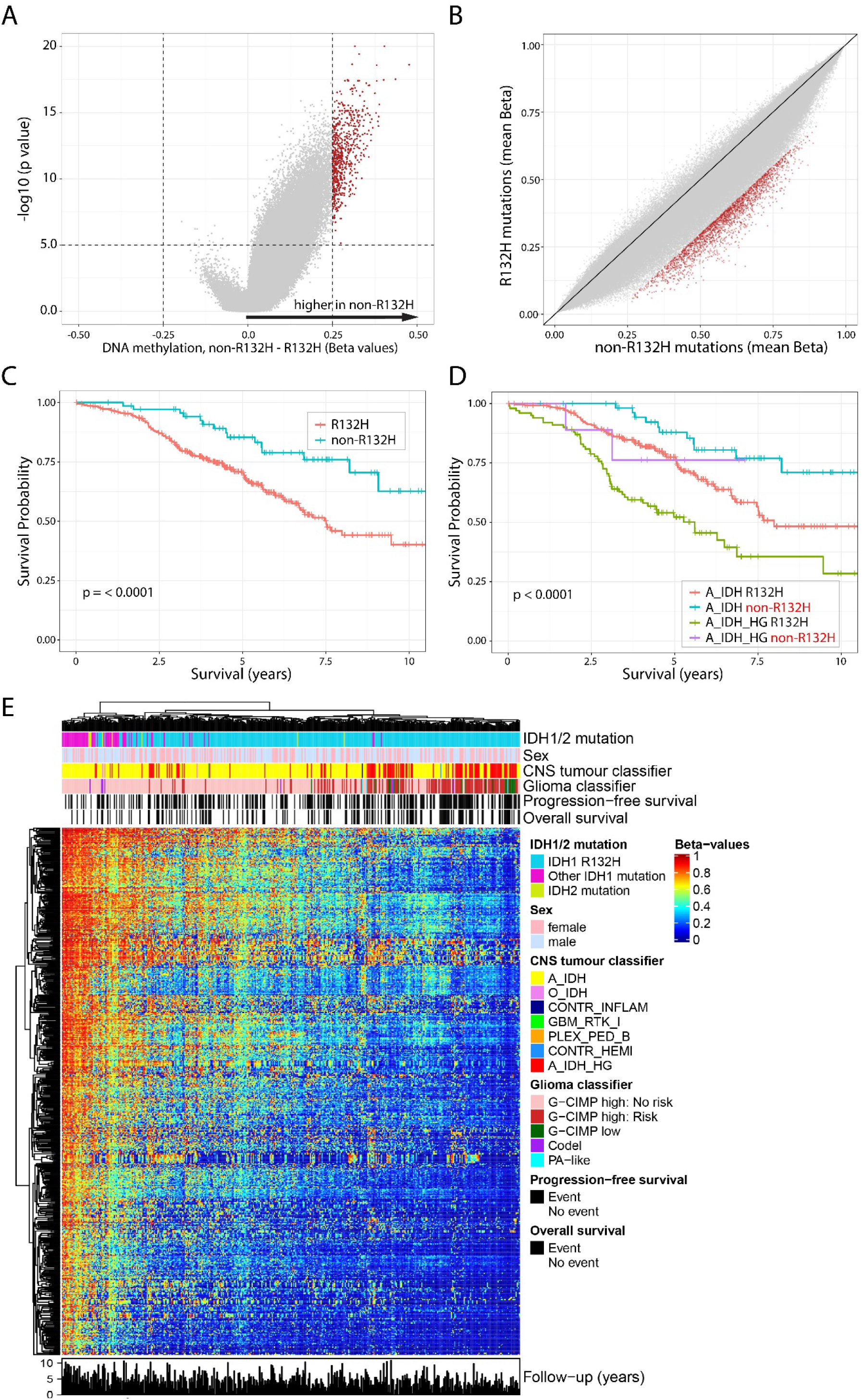
non-R132H mutations are associated with higher DNA methylation levels and improved survival of 1p19q non-codeleted astrocytoma patients included in the CATNON trial. Volcanoplot (A) and XY plot (B) showing differences in methylation in non-R132H v. IDH1^R132H^ mutated tumours. C) patients harbouring non-R132H mutated tumours have improved outcome, which is independent of methylation class (D). Heatmap of the most differentially methylated probes (red dots in A and B), shows a gradient in methylation levels. Non-R132H mutated tumours cluster at the far left (high methylation), where poor prognostic methylation subtypes (epigenetics subtypes) cluster at the opposite end.

Gliomas with higher levels of genome wide DNA methylation generally are associated with longer survival in adults^23,34–36^. Since non-R132H mutated gliomas have increased DNA methylation levels, we compared overall survival of patients with different IDH mutations. In patients included in the CATNON randomised phase III clinical trial, those harbouring tumours with non-R132H mutations indeed had longer overall survival compared to patients harbouring IDH1^R132H^ mutated tumours (figure 2C). The hazard ratio for non-R132H mutations was 0.41, 95% CI [0.24, 0.71], p=0.0013.

DNA methylation profiling can also assign tumours to specific (prognostic) methylation subclasses. In line with the poorer survival, IDH1^R132H^ mutated tumours also had a significantly higher proportion assigned to the prognostically poorer subclass A_IDH_HG (“IDH-mutant, high grade astrocytoma”, n=100/366 v. 9/71, p= 0.036, chi-squared test) using the subclasses as defined by Capper et al. (“CNS-classifier”)^37^. They also have a higher proportion of G-CIMP low tumours (18/369 v. 0/62) and G-CIMP-high tumours with risk to progression to G-CIMP low (111/335 v. 2/62) in the classifier as defined by the TCGA and de Souza et al. (“glioma classifier”, p< 0.001, chi-squared test, table 1) ^23,34^.

**Table 1.**
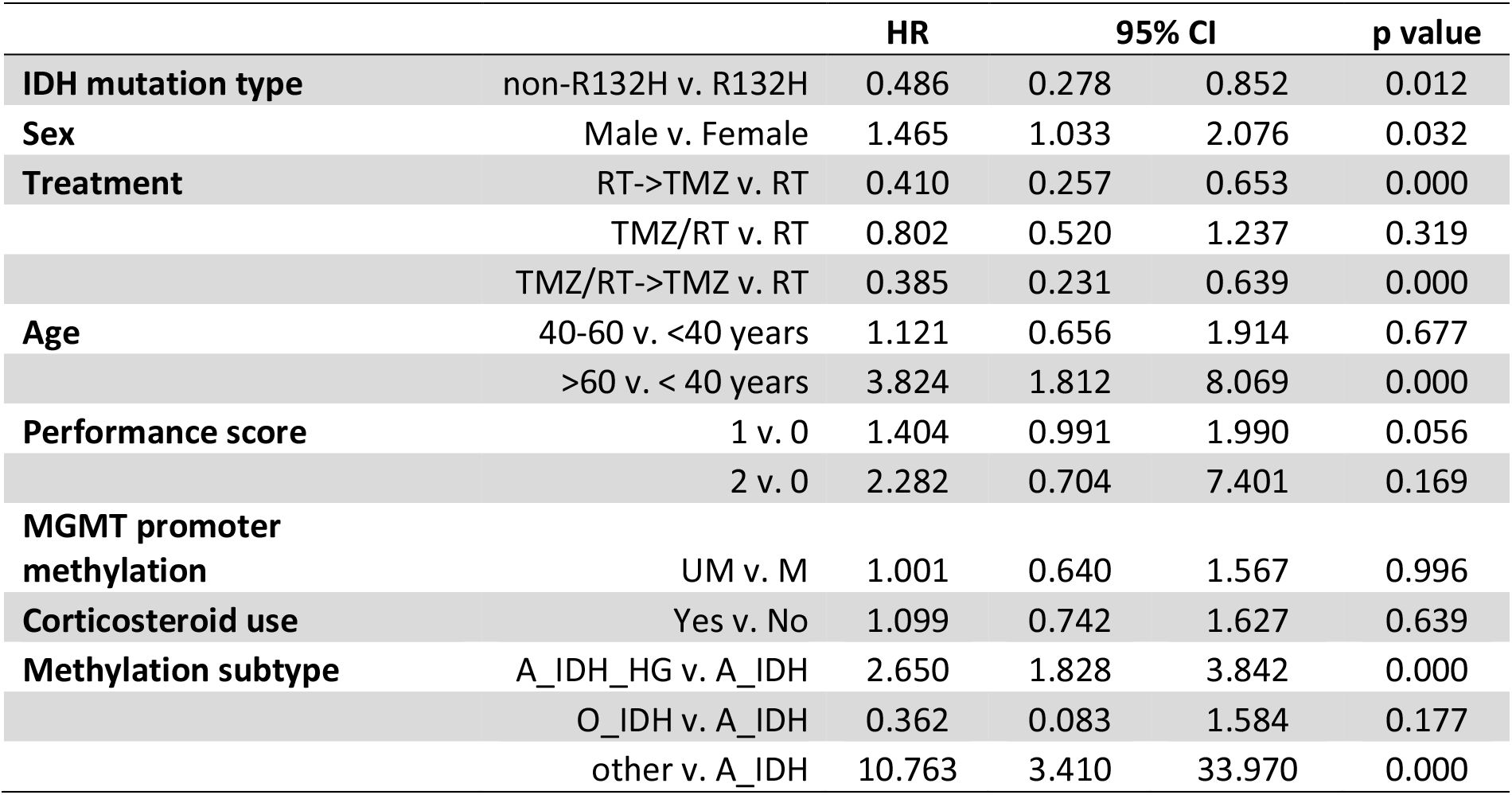
Multivariable model

|  |  | HR | 95% CI |  | p value |
| --- | --- | --- | --- | --- | --- |
| **IDH mutation type** | non-R132H v. R132H | 0.486 | 0.278 | 0.852 | 0.012 |
| **Sex** | Male v. Female | 1.465 | 1.033 | 2.076 | 0.032 |
| **Treatment** | RT->TMZ v. RT | 0.410 | 0.257 | 0.653 | 0.000 |
|  | TMZ/RT v. RT | 0.802 | 0.520 | 1.237 | 0.319 |
|  | TMZ/RT->TMZ v. RT | 0.385 | 0.231 | 0.639 | 0.000 |
| **Age** | 40-60 v. <40 years | 1.121 | 0.656 | 1.914 | 0.677 |
|  | >60 v. < 40 years | 3.824 | 1.812 | 8.069 | 0.000 |
| **Performance score** | 1 v. 0 | 1.404 | 0.991 | 1.990 | 0.056 |
|  | 2 v. 0 | 2.282 | 0.704 | 7.401 | 0.169 |
| **MGMT promoter methylation** | UM v. M | 1.001 | 0.640 | 1.567 | 0.996 |
| **Corticosteroid use** | Yes v. No | 1.099 | 0.742 | 1.627 | 0.639 |
| **Methylation subtype** | A_IDH_HG v. A_IDH | 2.650 | 1.828 | 3.842 | 0.000 |
|  | O_IDH v. A_IDH | 0.362 | 0.083 | 1.584 | 0.177 |
|  | other v. A_IDH | 10.763 | 3.410 | 33.970 | 0.000 |

A heatmap of the most differentially methylated CpGs of CATNON data (n=677, selected on a beta value change > 0.25 and false discovery corrected p values < 10e-5) shows a gradient from high to low methylation levels. As expected, the non-R132H mutated tumours cluster together at the high-methylation end of this spectrum. Interestingly, most of the tumours with less favourable molecular subtypes (A_IDH_HG, G-CIMP low, G-CIMP high with risk to progression) clustered together at the other, demethylated end (figure 2E). Although the clinical follow-up of CATNON patients is limited, the number of mortality events also tended to cluster at the demethylated end of the heatmap which suggests that there is a strong correlation between the level of methylation of these 677 probes and survival.

To determine whether the type of mutation is a prognostic factor independent of the DNA methylation subtypes, we stratified these subtypes by *IDH1/2* mutation (IDH1^R132H^ *v*. non-R132H). Our data show that, even within the prognostic DNA methylation subtypes, patients harbouring non-R132H mutated tumours had a significantly longer survival compared to those harbouring IDH1^R132H^-mutated tumours, regardless of the classifier used (figure 2D, supplementary figure 1). The type of *IDH1/2* mutation was also an independent prognostic factor in a multivariable analysis that included all known factors associated with survival in this trial (treatment, age, corticosteroid use and sex, supplementary table 1). It remained significant when DNA methylation subclass was included in this analysis (table 1, supplementary table 2). These data demonstrate that the type of *IDH1/2* mutation is an independent factor associated with patient survival.

To confirm these observations, we performed a similar analysis on the *IDH1/2* mutated, 1p/19q non-codeleted glioma patients included in the TCGA dataset^23^. Similar to observed in the CATNON dataset, a striking increase in DNA methylation levels was seen in non-R132H mutated tumours compared to those harbouring a IDH1^R132H^ mutation (figure 3AB). Also similar was the observation that patients harbouring non-R132H mutated tumours survived significantly longer; the HR of patients harbouring non-R132H mutated tumours (n=37) versus IDH1^R132H^-mutated tumours (n=177) was 0.20 (95% CI [0.047, 0.837], p=0.028 figure 3C). Finally, IDH1^R132H^ mutated tumours also had a higher proportion of tumours assigned to the prognostically poorer G-CIMP low DNA methylation class (4/116 v. 0/27) and a higher number at risk of progression to G-CIMP low (29/111 v. 0/24, p=0.016).

**Figure 3:**
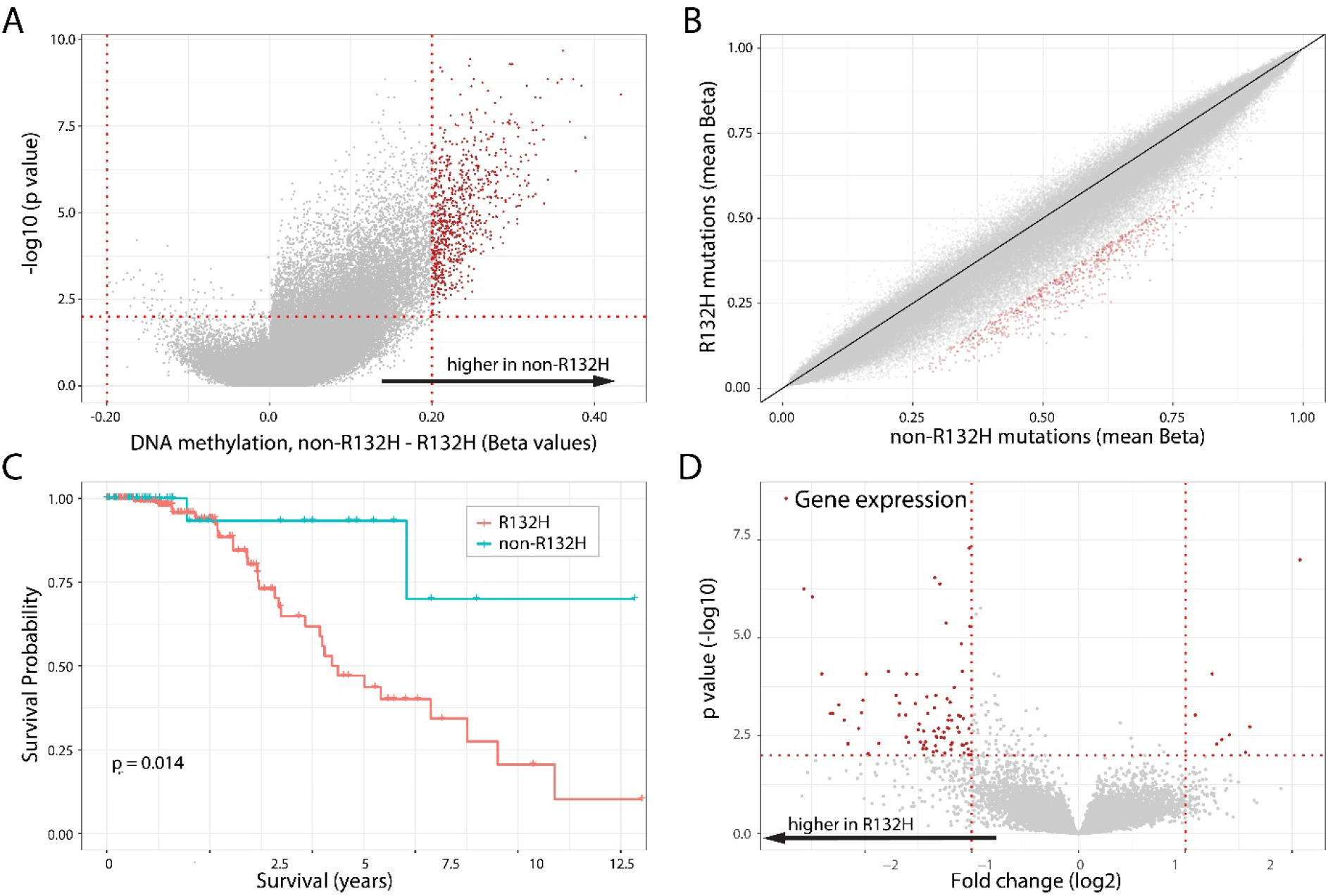
non-R132H mutations are associated with higher DNA methylation levels, lower gene expression and improved survival of 1p19q non-codeleted astrocytoma patients of the TCGA. Volcanoplot (A) and XY plot (B) showing differences in methylation in non-R132H v. IDH1^R132H^ mutated tumours. C) patients harbouring non-R132H mutated tumours have improved outcome. (D) Volcanoplot showing differential expresson of genes between non-R132H and IDH1^R132H^ mutated tumours. Most differentially expressed genes (red dots) have lower expression in non-R132H mutated tumours (see also supplementary table 2).

DNA methylation generally shows a negative correlation with gene expression, especially when the methylated CpGs are located near the transcriptional start site ^38,39^. We therefore examined whether the reduction in DNA methylation in IDH1^R132H^ mutated tumours is associated with an increase in gene expression in the 1p/19q non-codeleted gliomas present in the TCGA dataset. Indeed, of the genes differentially expressed between IDH mutation types (with > 2 fold change in expression level at p<0.01 significance level) in astrocytomas, most (157/183, 86%) were upregulated in IDH1^R132H^ mutated tumours (figure 3D, supplementary table 3). Pathway analysis using these 183 genes indicates that genes upregulated in IDH1^R132H^ mutated tumours were involved in cellular movement, cell death and survival, cell-to-cell signalling and interaction and carbohydrate metabolism (supplementary figure 2).

We performed a second validation using 1p/19q non-codeleted samples included in the randomised phase II TAVAREC clinical trial. Again, the vast majority of probes had lower DNA methylation levels in IDH1^R132H^ mutated tumours (n=83) compared to non-R132H mutated tumours (n=11, figure 4A) and the most differentially methylated probes were those partially methylated in IDH1^R132H^ mutated tumours (figure 4B). Moreover, there was a large degree of overlap in differential DNA methylation between CATNON and TAVAREC samples (figure 4C). In TAVAREC, there was no significant difference in survival between patients harbouring IDH1^R132H^ and non-R132H mutated tumours (HR 1.21, 95% CI [0.60, 2.45], P=0.60). This however, may be related to the specific inclusion criteria of this trial: patients were included only when the tumour showed signs of malignant progression at the time of progression (i.e. contrast enhancement on the MRI scan). In this respect it is interesting to note that the percentage of non-R132H mutated tumours was almost two-fold lower in TAVAREC trial samples (13%) compared to CATNON (19%) and TCGA (20%). Although this difference in frequency was not significant, these numbers are in line with the notion that non-R132H mutated tumours have lower frequencies of malignant progression. The small number of patients harbouring non-R132H mutated tumours (n=11) may also mask potential survival differences. A heatmap of most differentially methylated probes shows that non-R132H-mutated tumours and tumours assigned to the prognostically poorer subclass A_IDH_HG clustered at opposites ends of this heatmap (figure 4D). A forest plot of the combined CATNON, TCGA and TAVAREC survival data shows a summary estimate HR for non-R132H mutated tumours of 0.56 with 95% CI [0.37, 0.85], association p=0.006 (figure 4E).

**Figure 4:**
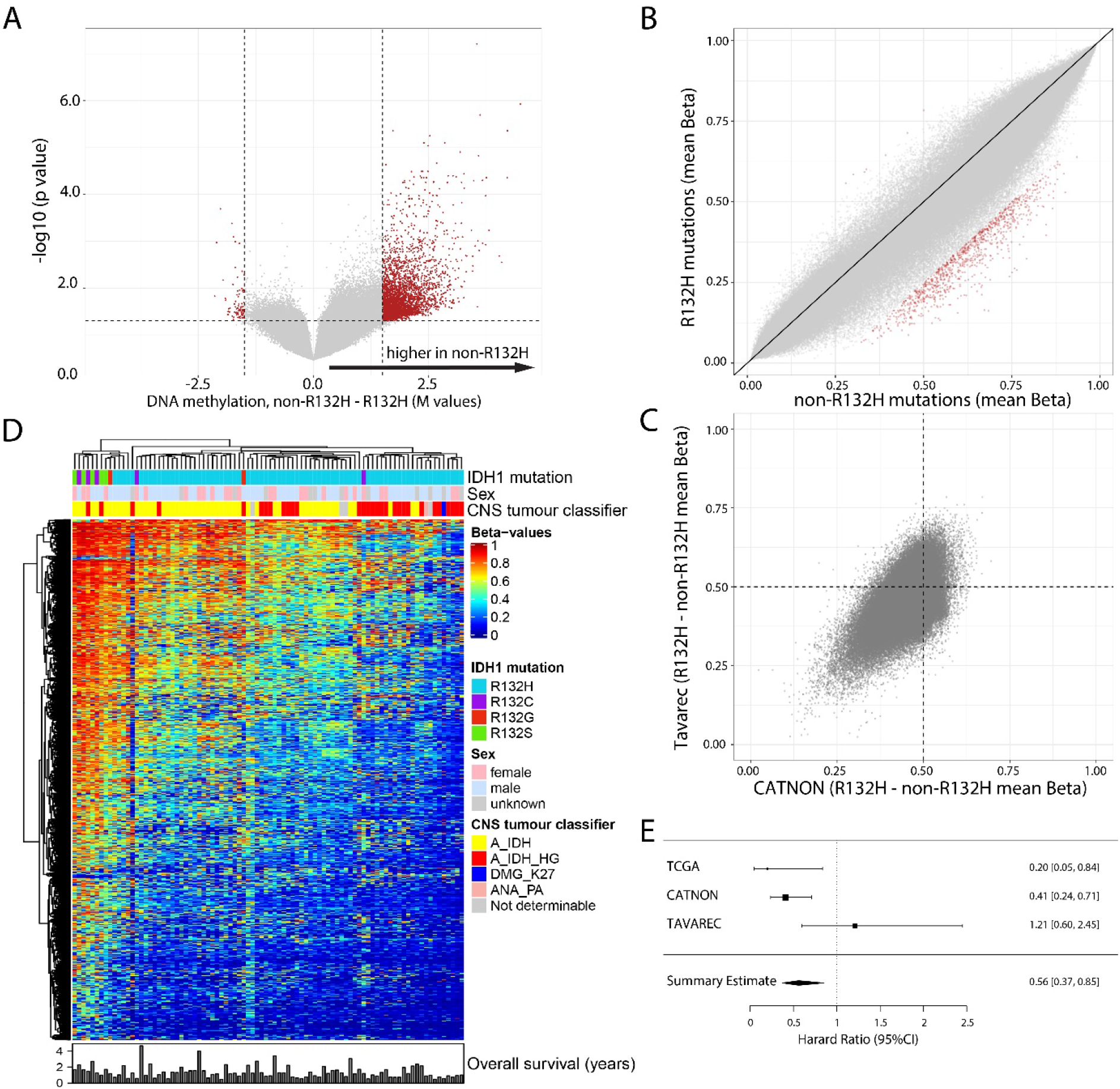
non-R132H mutations are associated with higher DNA methylation levels in 1p19q non-codeleted astrocytoma samples of patients included in the Tavarec trial. Volcanoplot (A) and XY plot (B) showing differences in methylation in non-R132H v. IDH1^R132H^ mutated tumours. C) Differential methylation between non-R132H v. IDH1^R132H^ mutated tumours showed a large degree of overlap in CATNON (x axis) and Tavarec (y axis) samples. (D) Heatmap of the most differentially methylated probes (red dots in A and B), shows a gradient in methylation levels. Non-R132H mutated tumours cluster at the far left (high methylation), where poor prognostic methylation subtypes (epigenetics subtypes) cluster at the opposite end. E) Forrest plot showing the summary HR estimate of 1p19q non-codeleted astrocytoma patients harbouring non-R132H v. IDH1^R132H^ mutated tumours.

To test whether mutation-dependent DNA methylation differences were restricted to 1p/19q non-codeleted gliomas, we analysed the genome-wide methylation profiles of i) *IDH1/2* mutated, 1p/19q codeleted gliomas (TCGA) ii) acute myeloid leukemias (TCGA) and iii) chondrosarcomas. Although the sample sizes of these datasets were relatively small in all tumour types (1p/19q codeleted gliomas n=135 *v*. 14; acute myeloid leukemias n=4 *v*. n=24; chondrosarcomas n=3 *v*. n=17 for IDH1^R132H^ and non-R132H mutated tumours respectively), there was less DNA methylation in IDH1^R132H^ *v*. non-R132H mutation tumours (figure 5A-C). These data demonstrate that the level of DNA methylation is lower in tumours harbouring *IDH1/2* mutations with presumed low D-2HG production.

**Figure 5:**
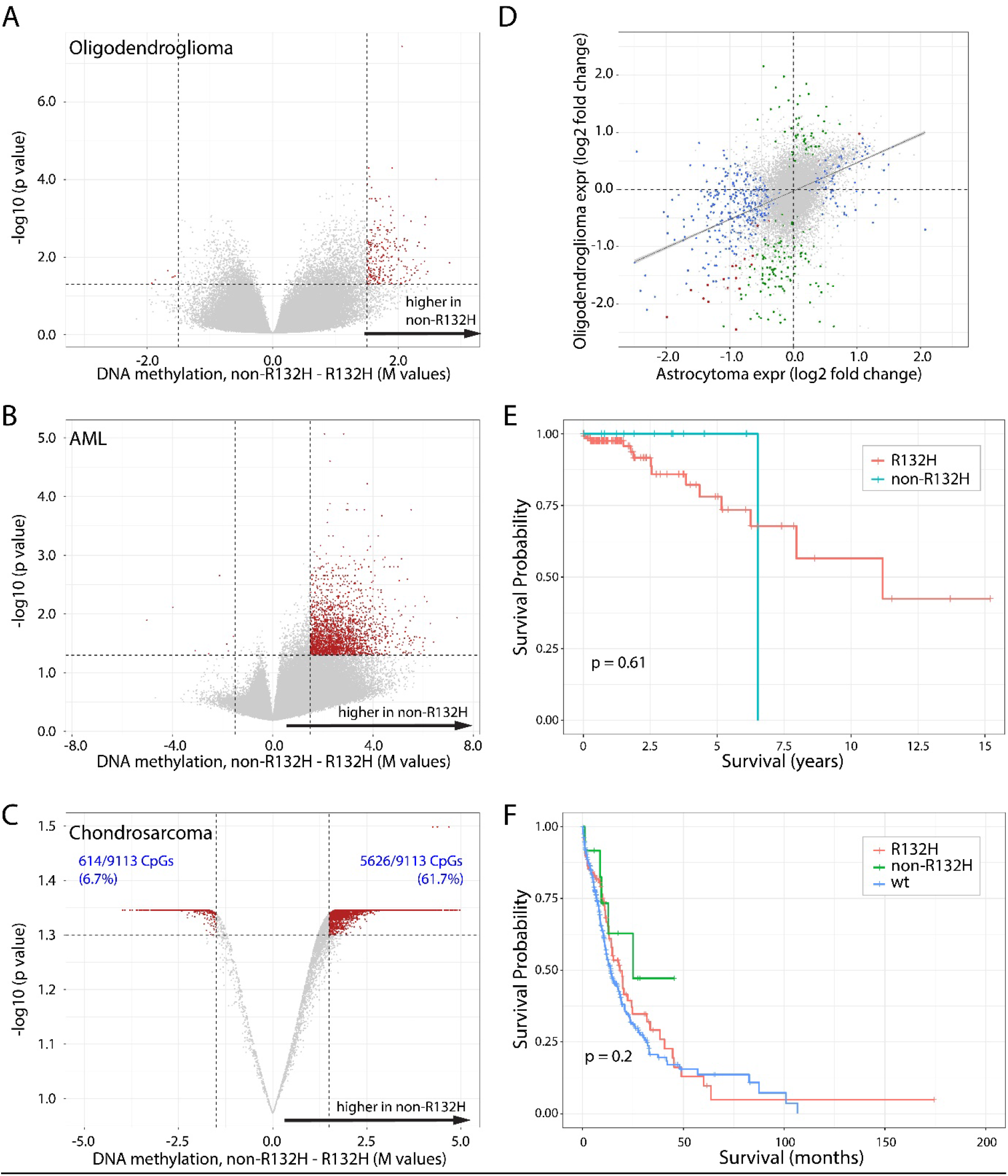
non-R132H mutations are associated with higher DNA methylation levels independent of tumour type. Volcanoplot of 1p19q codeleted oligodendrogliomas (A), AML (B) and chondrosarcomas (C) showing differences in methylation in non-R132H v. IDH1^R132H^ mutated tumours. Red dots depict CpGs that had a > 0.2 change in beta value, and were significant (P<0.01) in a Wilcoxon rank test (in B and C, the Y-axis are t-test P-values for visualization purposes). Although the difference in chondrosarcomas is less than in other tumour types, the majority of significant CpGs was in non-R132H-mutated tumours (e.g. 225 CpG showed a > 0.3 increase in beta value at p<0.01 where only 47 showed a similar decrease). D) Gene expression differences between non-R132H v. IDH1^R132H^ mutated tumours in 1p19q non-codeleted astrocytomas (x-axis) and 1p19q codeleted oligodendrogliomas (y-axis) shows a large degree of overlap. Blue, green and red dots depict genes significantly differentially expressed in astrocytomas, oligodendrogliomas or both respectively (see also supplementary table 2 and 3). E). Survival of 1p19q codeleted oligodendroglioma patients present in the TCGA database harbouring non-R132H v. IDH1^R132H^ mutated tumours. There were too few events evaluate survival differences per mutation type. F) mutation type-specific survival differences in AML.

Gene expression analysis of 1p/19q codeleted gliomas present in the TCGA dataset identified 148 differentially expressed genes (expression fold change >1 or <-1 and p< 0.01). Similar to observed in astrocytic tumours, the majority of identified genes (123/148, 83%) were upregulated in IDH1^R132H^ mutated tumours (supplementary table 4). Moreover, there was a relatively large degree of concordance in differential expression between the two analyses (figure 5D) and sixteen genes were identified in both analyses.

The number of samples and events of the various datasets in patients with 1p/19q codeleted gliomas was insufficient to determine mutation type dependent survival differences. For example, there were only 14 non-R132H mutated 1p/19q codeleted tumours in the TCGA dataset, with only 1 event noted (in the IDH1^R132H^ mutated tumours there were 14 events in 135 patients). The HR for TCGA samples was 0.59 (95% CI [0.077, 4.595], p= 0.62, figure 5E). Also in the MSK-Impact^26^ and the Chinese Glioma Genome Atlas (CGGA)^29^ there were too few samples and events to determine survival benefit in patients harbouring non-R132H-mutated tumours. In these datasets, the events/number in non-R132H *v*. IDH1^R132H^ mutated samples was 0/6 *v*. 3/34 and 0/5 *v*. 3/31 in MSK impact, and CGGA datasets respectively. We were not able to determine survival differences in AML (n=12 with 5 events *v*. n=89, 54 events, HR 1.49, 95%CI[0.59, 3.75], p=0.39, figure 5F).

## Discussion

Our data shows that *IDH1/2mt* gliomas are distinct when compared to other *IDH1/2mt* tumours in that they have a disproportionally high percentage of IDH1^R132H^ mutations and raise the attractive clinical association between different rarer (codon 132) mutations and outcome. Patients harbouring IDH1^R132H^ mutated tumours have lower levels of genome-wide DNA methylation, regardless of tumour type (1p/19q non-codeleted gliomas, 1p/19q codeleted gliomas, AML and chondrosarcomas). For 1p/19q non-codeleted *IDH1/2mt* gliomas, this difference is clinically relevant as patients harbouring non-R132H mutated tumours have improved outcome. Since IDH1^R132H^ mutations are presumed to be relatively poor in D-2HG production, our data are in line with the observation that glioma patients with higher D-2HG levels have improved outcome^40^. Our data are also in line with data from a meeting abstract showing similar mutation-specific survival differences^41^

The observation that patients harbouring non-R132H mutated gliomas have longer survival is of importance for clinical practice as the specific *IDH1/2* mutation could alter patient prognostication. In this respect diagnostic assays should be able to discriminate between the type of IDH-mutation present; non-R132H mutations comprise of up to 20% of all IDH-mutations in 1p/19q non-codeleted gliomas. Moreover, the efficacy of treatment with alkylating agents, IDH1/2 inhibitors, or other novel treatments might vary per mutation type, and therefore may be taken into account as stratification factor in future clinical trials.

It has been reported that D-2HG is a relatively weak inhibitor of TET2. In fact, the IC50 value for TET2 inhibition (~5 mM) is in the same range as the intratumoural D-2HG levels^18,33,42,43^. As TET2 mediates the first step in DNA demethylation, lower D-2HG levels may result in reduced inhibition of DNA-demethylation. Such lower D-2HG levels have been reported for IDH1^R132H^ mutated tumours in some studies ^18,43,44^ (but not in all ^42^), though confounding factors such as tumour purity may influence these observations. In addition, the *K*_M_ for D-2HG production of the IDH1^R132H^ mutation is higher than that of other IDH1 mutations (though the enzymatic efficiency may be similar for some mutations) ^17^. Although we did not directly measure D-2HG levels, the partial inhibition of TET2, may explain the lower overall methylation in IDH1^R132H^-mutated tumours.

The improved outcome of non-R132H mutated astrocytomas may be explained by a reduced expression of genes that support tumour growth and/or induce treatment sensitivity caused by the increase in CpG methylation. Evidence supporting this hypothesis is the observation that many of the differentially expressed genes are involved in pathways associated with cancer. However, the improved outcome of non-R132H mutated astrocytomas may also be related to the observation that D-2HG is toxic to cells, though only at high concentrations. For example, we have previously shown that exposure to D-2HG or expression of mutated IDH constructs reduced proliferation of cells, both *in-vitro* and *in-vivo*^45^. Later independent studies largely confirmed these observations and also conversely, reduction of D-2HG levels by mutant IDH inhibitors increased cell proliferation^16,46–49^. It should be noted however, that in some preclinical model systems a growth inhibitory effect of IDH-inhibitors was observed^50,51^. Functional experiments should confirm this hypothesis. Alternatively, differences in genetic stress and related mutational signatures may also explain the differential distribution of mutations in IDH^32,52^.

Limitations of this study include the relatively small sample size of several datasets, especially those with diagnosis other than the non-1p/19q codeleted gliomas. In addition, the absence of D-2HG level data limits the exploration of a direct correlation between *IDH1/2* mutation type and genome-wide DNA methylation.

In short, we described the effect of *IDH1/2* mutation type on patient outcome and the strong correlation between these specific mutations and genome-wide DNA methylation status. Our observation that non-R132H-mutated 1p/19q non-codeleted gliomas have a more favourable prognosis than their IDH1^R132H^ mutated counterpart is clinically relevant and should be taken into account for patient prognostication.

## Acknowledgements

The CATNON study was funded by Merck, Sharp and Dohme, and the Brain Tumor Group. The genome-wide DNA methylation profiles study was funded by grant GN-000577 from The Brain Tumour Charity, grant 10685 from the Dutch Cancer Society, financial support from the Vereniging Heino ‘Strijd van Salland’, and grant CA170278 from the United States Department of Defence. The authors thank the European Organization for Research and Treatment of Cancer for permission to use the data from EORTC studies 26053/22054 (CATNON) and 26091 (TAVAREC) for this research.

## Author contributions

Conceptualization: PJF

Methodology: CMT, YH, PJF

Validation: CMT

Investigation CMT, WRV, IdH, MdW, LB, PJF

Resources: MS, WT, PMC, WW, AAB, JFV, OLC, HW, SG, MG, LR, RR, MW, CMcB, JR, RHE, FC, TL, SC, AG, EL, FdV, PJM, MJBT, SV, JO, JVMGB, SE, MAV, AKN, WPM, JMK, PW, KA, RBJ, HJD, BB, VG, MvdB Data curation CMT, TG, PJF

Writing-original draft: CMT, WRV, MvdB, PJF

Writing-review and editing: all authors

Visualization: CMT, PJF Supervision: MvdB, PJF

## Competing interests

MS reports research grants from Astra-Zeneca, travel grant from Abbvie, personal fees from Genenta, outside the submitted work, PM reports support to attend conferences from BMS and an award towards an investigator initiated study from BMS. BB reports a MERCK grant for the EORTC22033 lGG study. MAV has indirect equity interest and royalty rights from Infuseon Therapeutics, Inc. He has received honoraria from Tocagen, Cellinta, and Celgene. None of these interests overlap with the research presented in this manuscript. Wolfgang Wick receives trial funding from Apogenix, Boehringer Ingelheim, Pfizer and Roche to the institution. He serves on advisory boards for Agios, Bayer, MSD, Novartis, Roche with compensation paid to the institution. MJvdB reports grants from Dutch Cancer Foundation, grants from Brain Tumor Charity, grants from Strijd van Salland, grants from MSD formerly Schering Plough, during the conduct of the study; personal fees from Carthera, personal fees from Nerviano, personal fees from Bayer, personal fees from Celgene, personal fees from Agios, personal fees from Abbvie, personal fees from Karyopharm, personal fees from Boston Pharmaceuticals, personal fees from Genenta, outside the submitted work. AN received research funding from Astra Zeneca, and Douglas Pharmaceuticals, consultancies for Bayer, Roche, Boehringer Ingelheim, MSD, Douglas Pharmaceuticals, Pharmabcine, Atara biotherapeutics, Trizell and Seagen. MW has received research grants from Abbvie, Adastra, Merck, Sharp & Dohme (MSD), Merck (EMD), Novocure and Quercis, and honoraria for lectures or advisory board participation or consulting from Abbvie, Adastra, Basilea, Bristol Meyer Squibb (BMS), Celgene, Medac, Merck, Sharp & Dohme (MSD), Merck (EMD), Nerviano Medical Sciences, Novartis, Orbus, Philogen, Roche and Tocagen. FdV reports support from AbbVie, Bioclin Therapeutics, BMS, GSK, Novartis, Octimed Oncology and Vaximm, outside the submitted work. PC reports support from BMS, AbbVie, Merck Serono, MSD, Vifor Pharma, Daiichi Sankyo, Leo Pharma and Astra Zeneca, ouside the submitted work. PJF reports grants from Dutch Cancer Foundation, the Brain Tumor Charity, the Strijd van Salland, de Westlandse ride and Hersentumorfonds, outside submitted work. Other authors report no conflict of interest.

---

**Supplementary figure 1:**
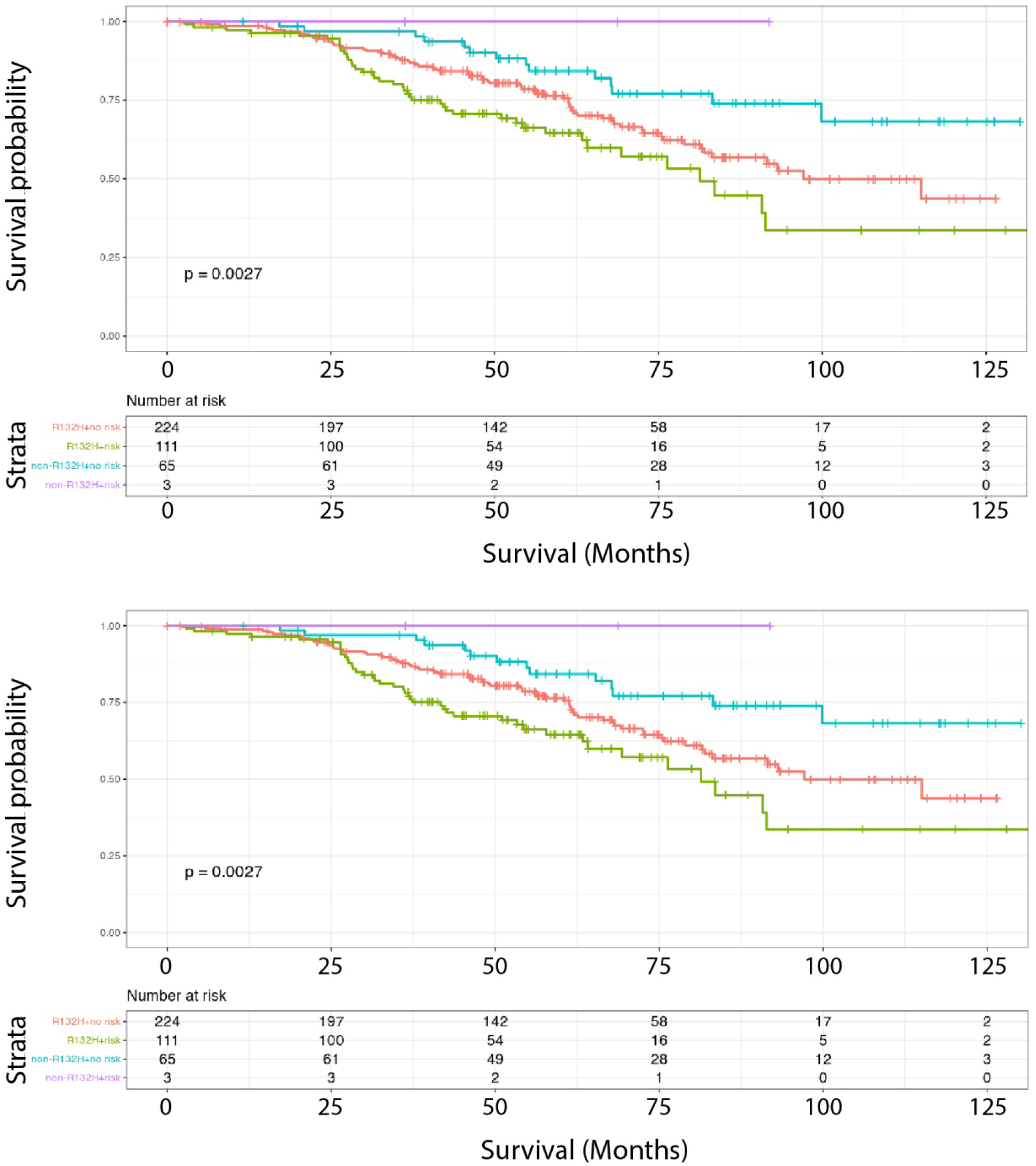
non-R132H mutations are associated with improved survival of 1p19q non-codeleted astrocytoma patients included in the CATNON trial independent of methylation class. Survival of patients harbouring non-R132H or IDH1^R132H^ mutated tumours stratified by methylation class as defined by the TCGA (top) or risk to G-CIMP-low progression (bottom). As can be seen, patients harbouring non-R132H mutated tumours have improved outcome, independent of methylation class.

**Supplementary figure 2:**
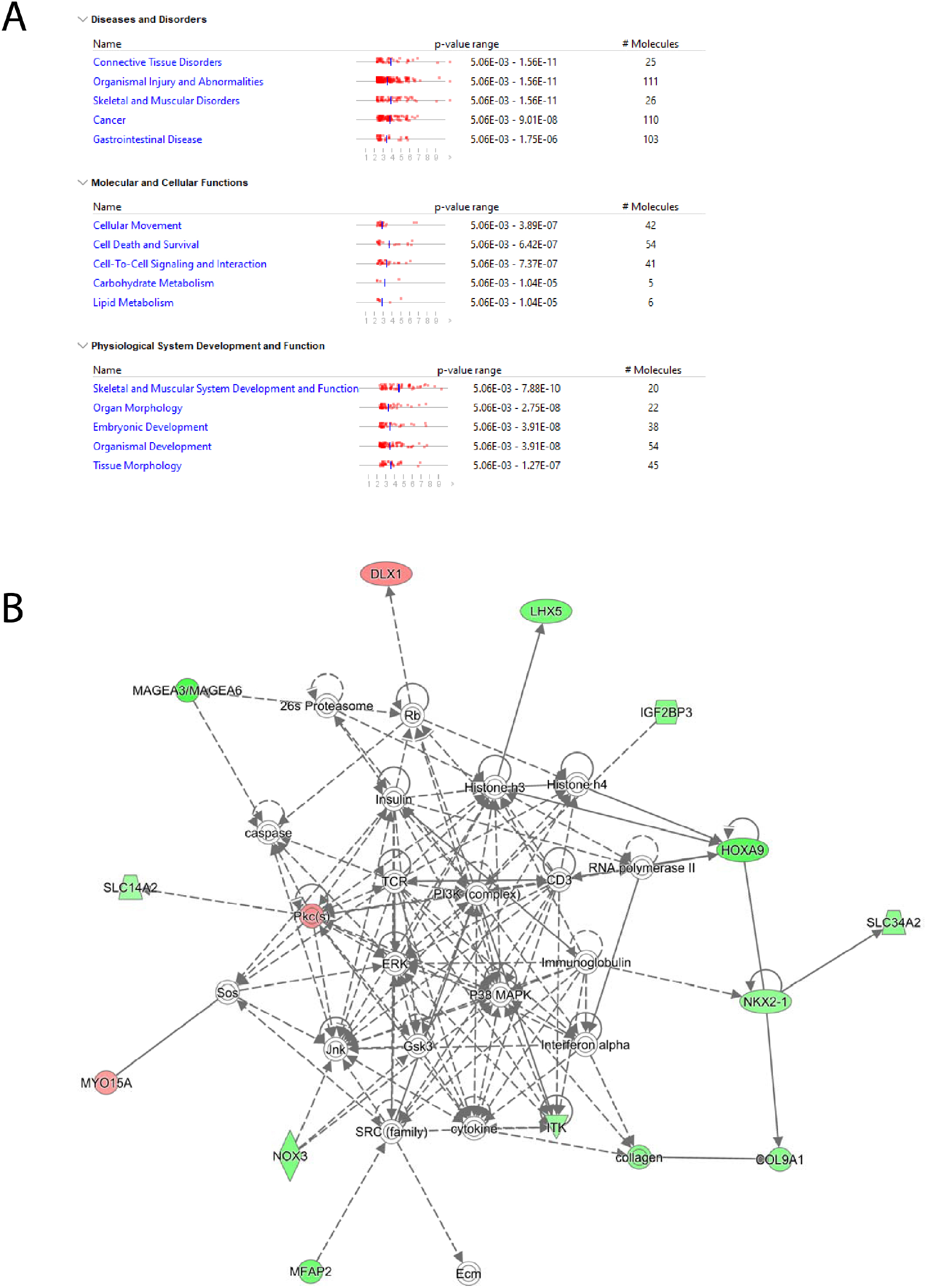
Pathway analysis of genes differentially expressed between non-R132H and IDH1^R132H^ mutated tumours. A) top diseases and disorders, molecular and cellular functions and physiological system development and function identified by pathway analysis. B) graphical representation of the top cancer pathway.

**Supplementary table 1.**
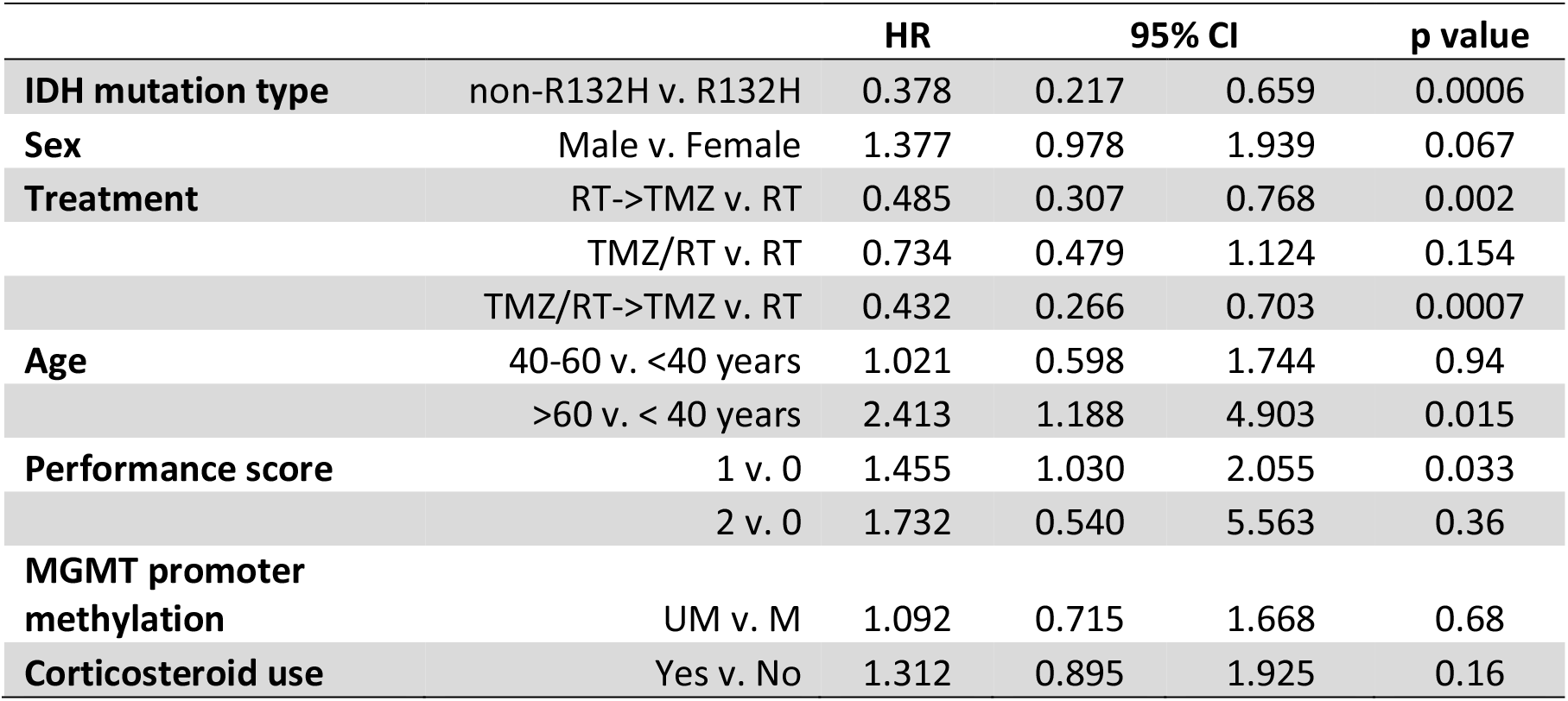

**Supplementary table 2.**
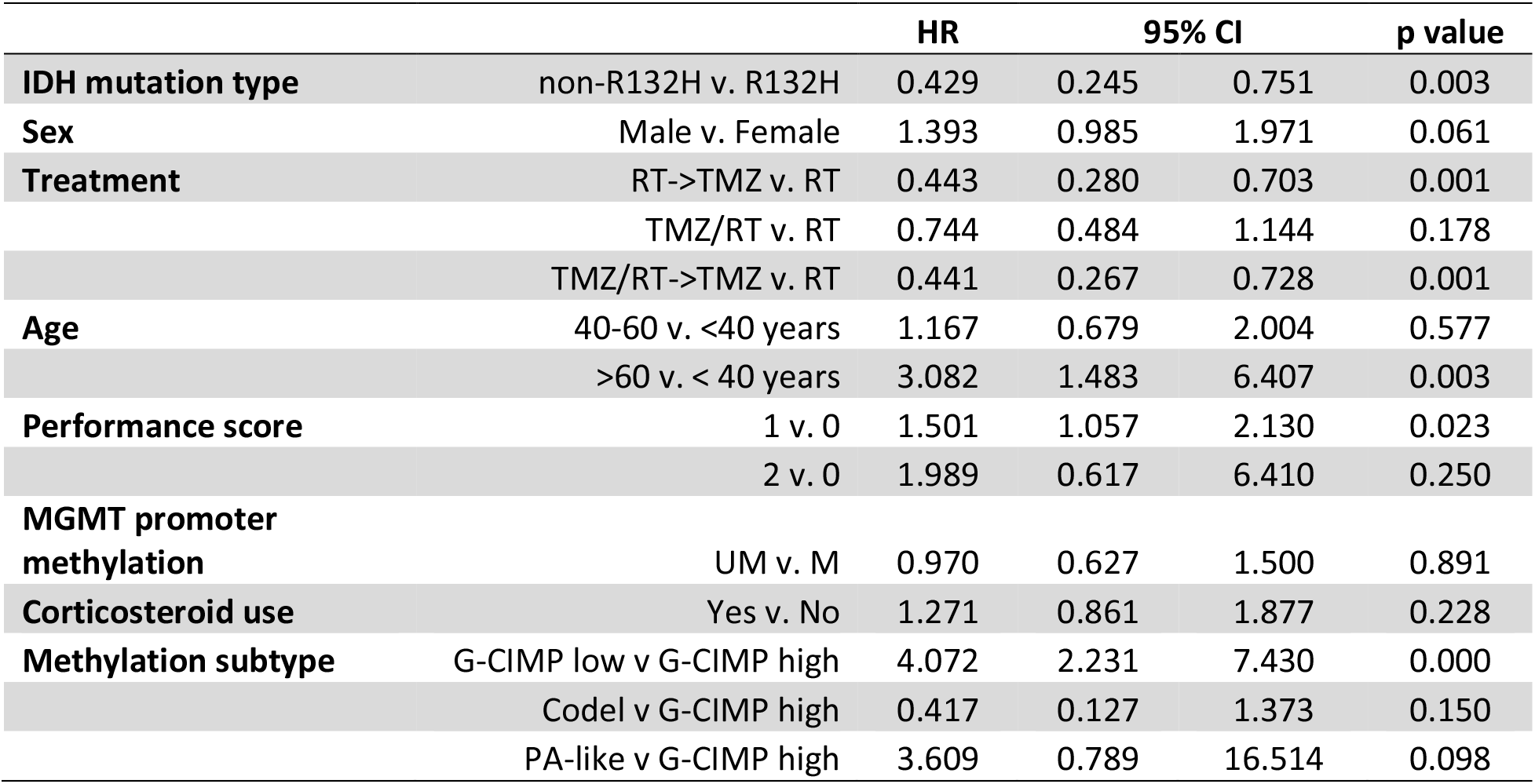

**Supplementary table 3.**
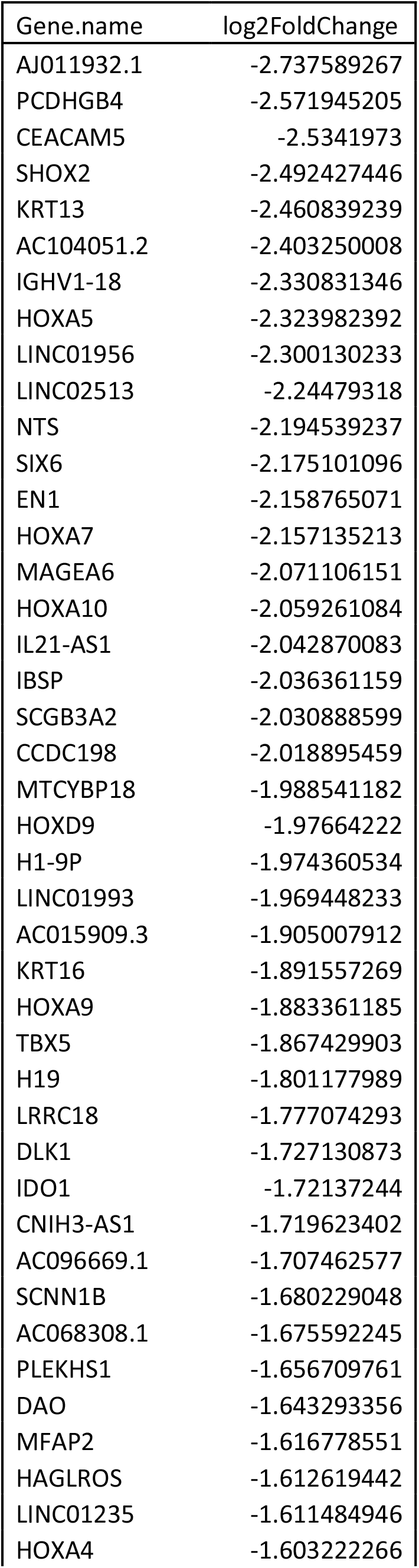
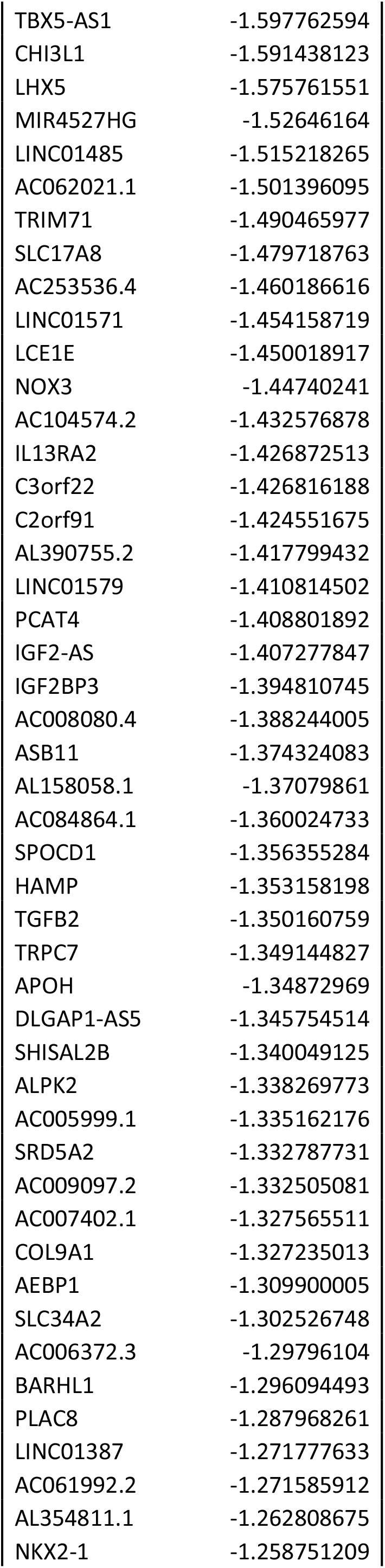
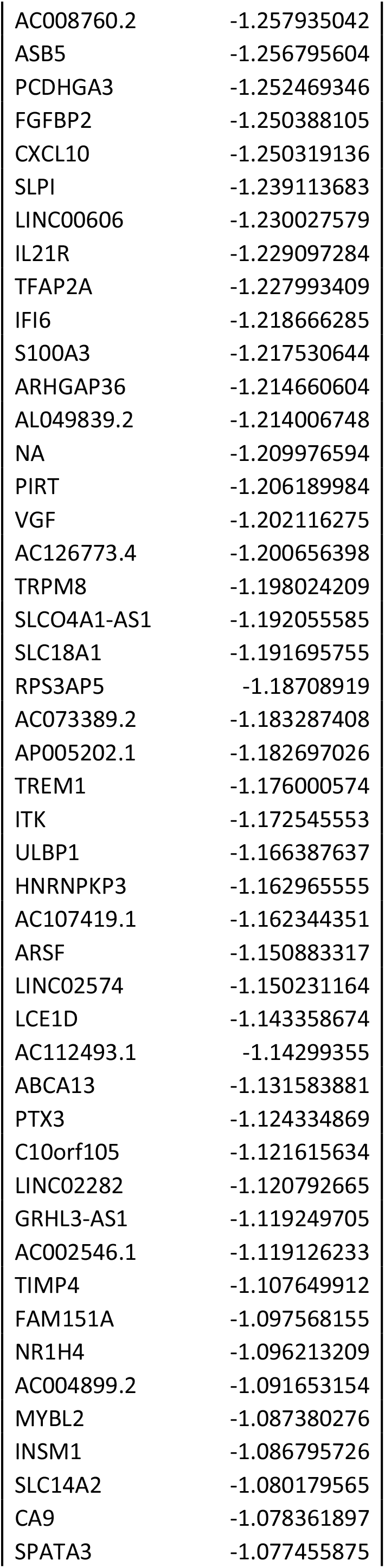
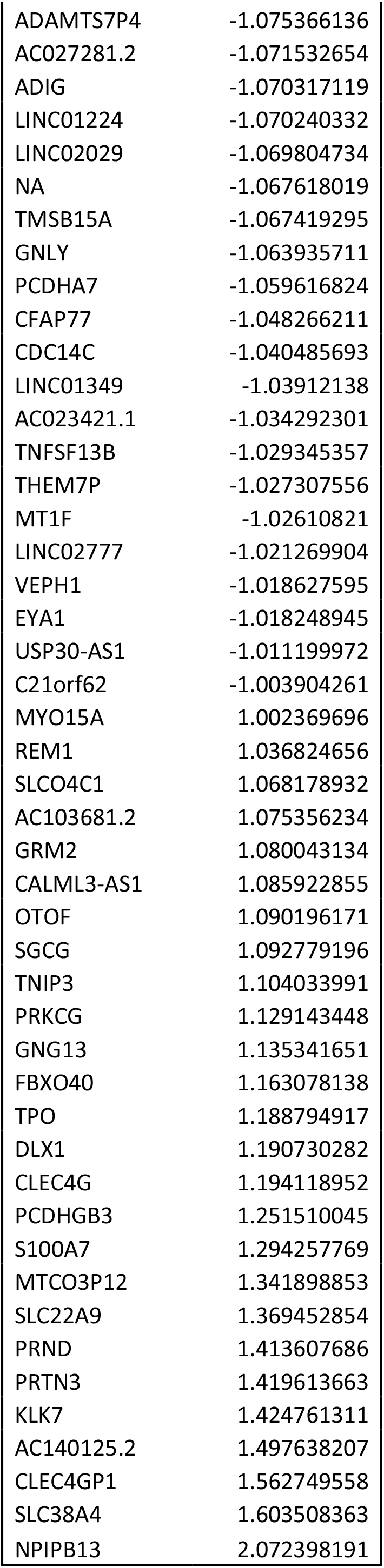

**Supplementary table 4.**
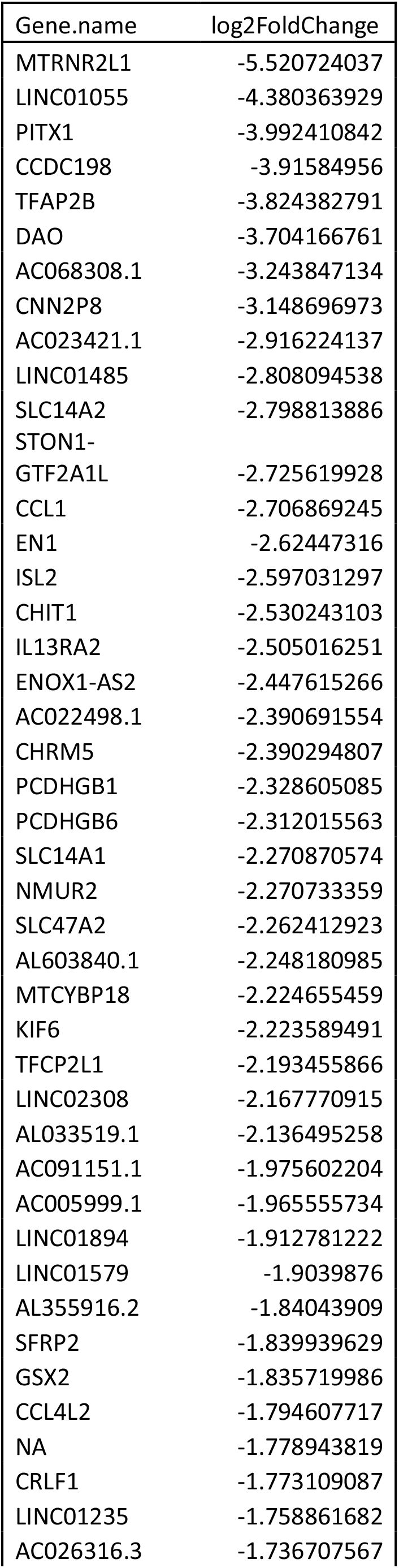
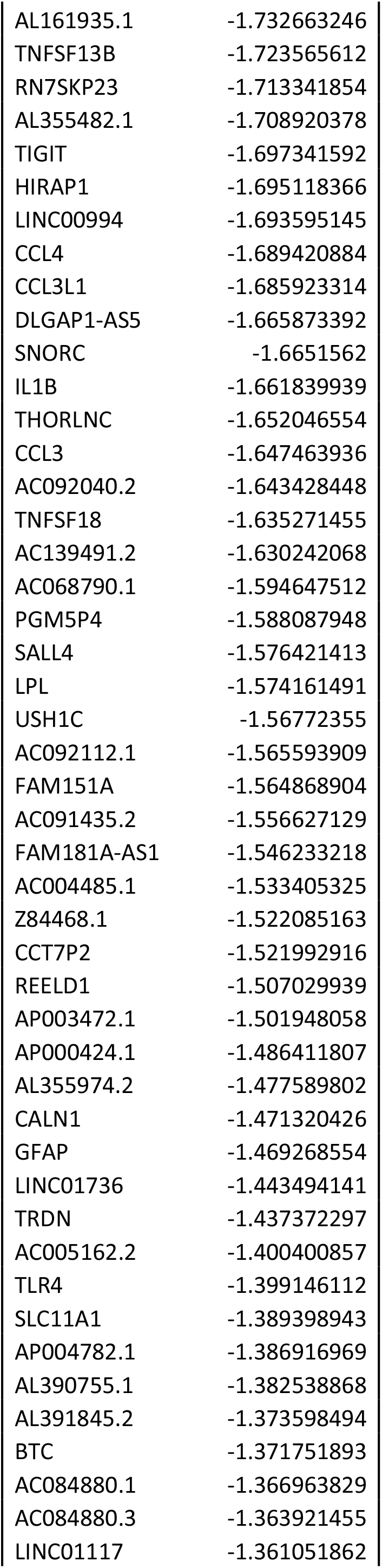
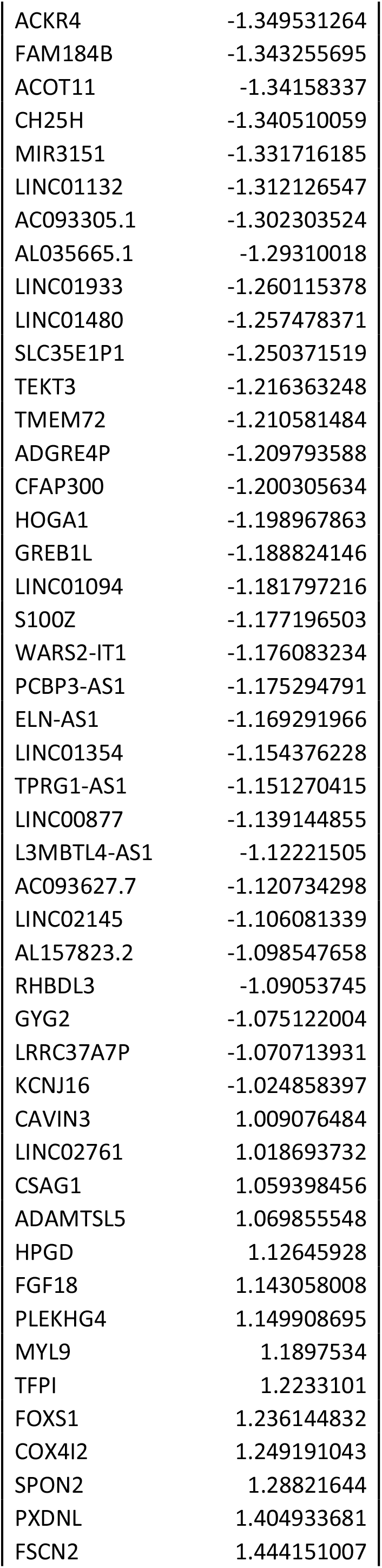
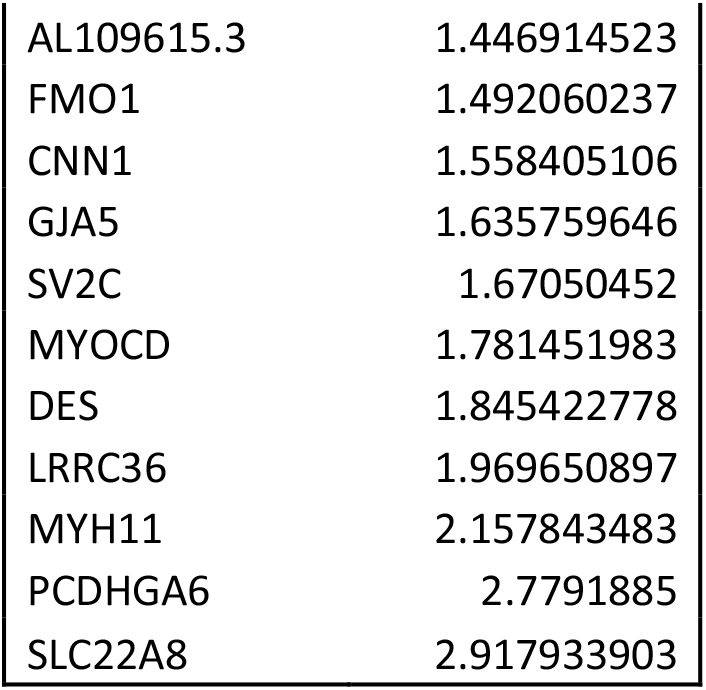

