## Supplementary figures and tables for "DNA methylation and survival differences associated with the type of IDH mutation in 1p/19q non-codeleted astrocytomas"

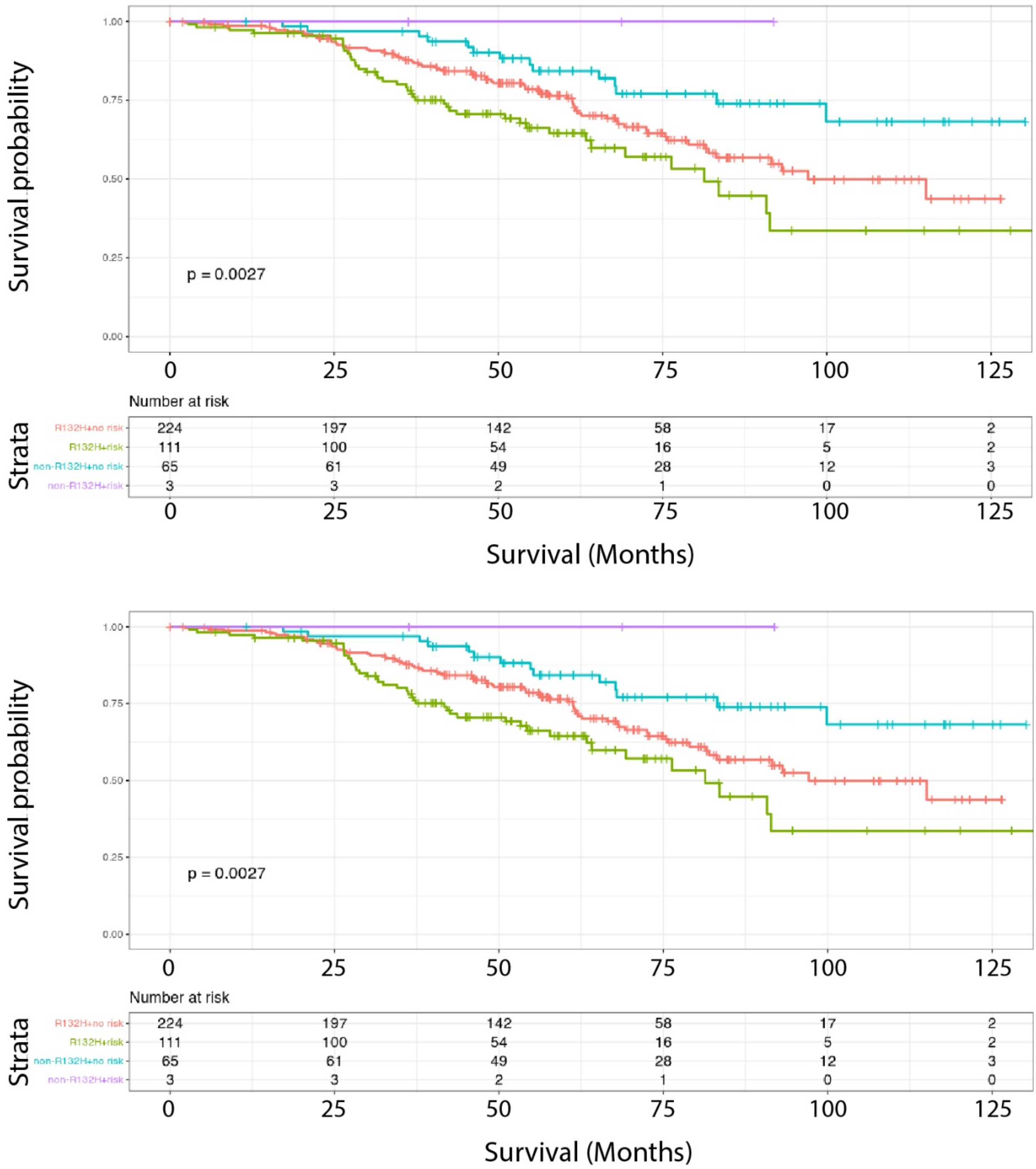


<u>Supplementary figure 1: non-R132H mutations are associated with improved survival of 1p19q non-codeleted astrocytoma patients included in the CATNON trial independent of methylation class</u>. Survival of patients harbouring non-R132H or IDH1$^{R132H}$ mutated tumours stratified by methylation class as defined by the TCGA (top) or risk to G-CIMP-low progression (bottom). As can be seen, patients harbouring non-R132H mutated tumours have improved outcome, independent of methylation class.

A

**Diseases and Disorders**

| Name | p-value range | # Molecules |
|---|---|---|
| Connective Tissue Disorders | 5.06E-03 - 1.56E-11 | 25 |
| Organismal Injury and Abnormalities | 5.06E-03 - 1.56E-11 | 111 |
| Skeletal and Muscular Disorders | 5.06E-03 - 1.56E-11 | 26 |
| Cancer | 5.06E-03 - 9.01E-08 | 110 |
| Gastrointestinal Disease | 5.06E-03 - 1.75E-06 | 103 |

**Molecular and Cellular Functions**

| Name | p-value range | # Molecules |
|---|---|---|
| Cellular Movement | 5.06E-03 - 3.89E-07 | 42 |
| Cell Death and Survival | 5.06E-03 - 6.42E-07 | 54 |
| Cell-To-Cell Signaling and Interaction | 5.06E-03 - 7.37E-07 | 41 |
| Carbohydrate Metabolism | 5.06E-03 - 1.04E-05 | 5 |
| Lipid Metabolism | 5.06E-03 - 1.04E-05 | 6 |

**Physiological System Development and Function**

| Name | p-value range | # Molecules |
|---|---|---|
| Skeletal and Muscular System Development and Function | 5.06E-03 - 7.88E-10 | 20 |
| Organ Morphology | 5.06E-03 - 2.75E-08 | 22 |
| Embryonic Development | 5.06E-03 - 3.91E-08 | 38 |
| Organismal Development | 5.06E-03 - 3.91E-08 | 54 |
| Tissue Morphology | 5.06E-03 - 1.27E-07 | 45 |

B

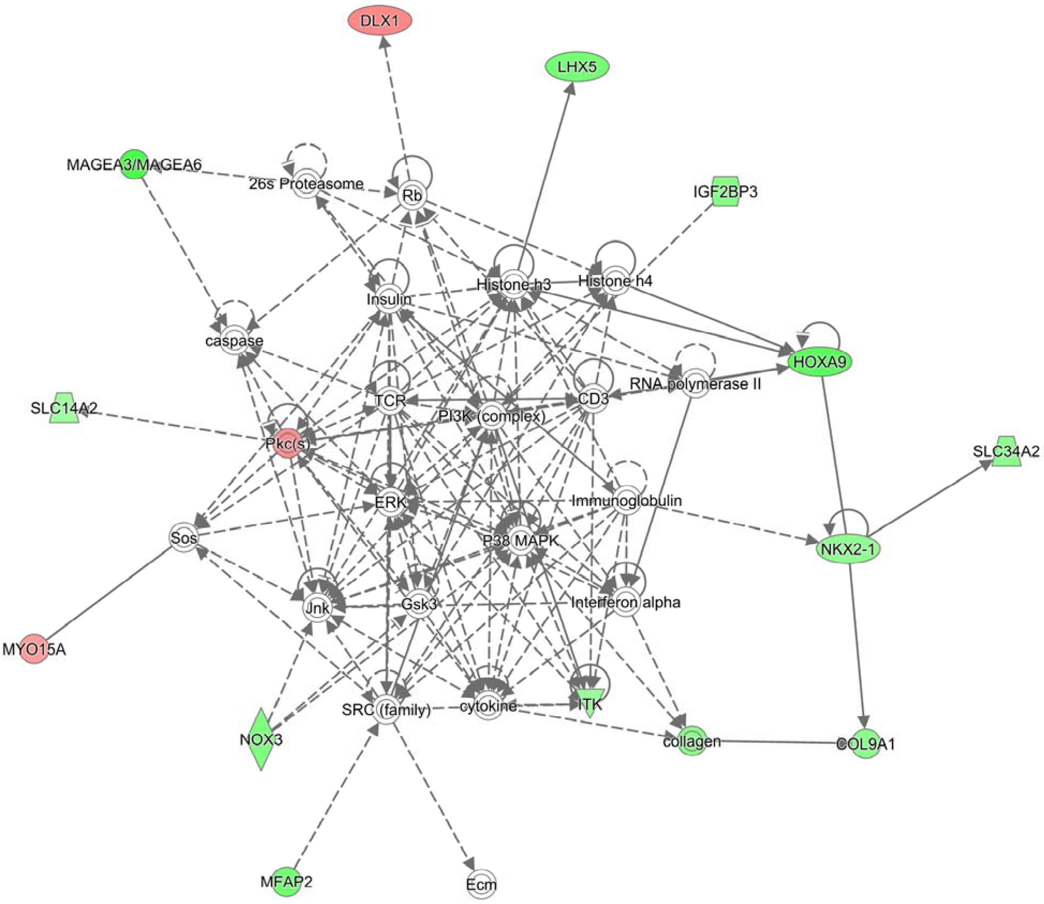


<u>Supplementary figure 2: Pathway analysis of genes differentially expressed between non-R132H and IDH1$^{R132H}$ mutated tumours.</u> A) top diseases and disorders, molecular and cellular functions and physiological system development and function identified by pathway analysis. B) graphical representation of the top cancer pathway.

Supplementary table 1

| | | HR | 95% CI | | p value |
|---|---|---|---|---|---|
| **IDH mutation type** | non-R132H v. R132H | 0.378 | 0.217 | 0.659 | 0.0006 |
| **Sex** | Male v. Female | 1.377 | 0.978 | 1.939 | 0.067 |
| **Treatment** | RT->TMZ v. RT | 0.485 | 0.307 | 0.768 | 0.002 |
| | TMZ/RT v. RT | 0.734 | 0.479 | 1.124 | 0.154 |
| | TMZ/RT->TMZ v. RT | 0.432 | 0.266 | 0.703 | 0.0007 |
| **Age** | 40-60 v. <40 years | 1.021 | 0.598 | 1.744 | 0.94 |
| | >60 v. < 40 years | 2.413 | 1.188 | 4.903 | 0.015 |
| **Performance score** | 1 v. 0 | 1.455 | 1.030 | 2.055 | 0.033 |
| | 2 v. 0 | 1.732 | 0.540 | 5.563 | 0.36 |
| **MGMT promoter methylation** | UM v. M | 1.092 | 0.715 | 1.668 | 0.68 |
| **Corticosteroid use** | Yes v. No | 1.312 | 0.895 | 1.925 | 0.16 |

Supplementary table 2

| | | HR | 95% CI | | p value |
|---|---|---|---|---|---|
| **IDH mutation type** | non-R132H v. R132H | 0.429 | 0.245 | 0.751 | 0.003 |
| **Sex** | Male v. Female | 1.393 | 0.985 | 1.971 | 0.061 |
| **Treatment** | RT->TMZ v. RT | 0.443 | 0.280 | 0.703 | 0.001 |
| | TMZ/RT v. RT | 0.744 | 0.484 | 1.144 | 0.178 |
| | TMZ/RT->TMZ v. RT | 0.441 | 0.267 | 0.728 | 0.001 |
| **Age** | 40-60 v. <40 years | 1.167 | 0.679 | 2.004 | 0.577 |
| | >60 v. < 40 years | 3.082 | 1.483 | 6.407 | 0.003 |
| **Performance score** | 1 v. 0 | 1.501 | 1.057 | 2.130 | 0.023 |
| | 2 v. 0 | 1.989 | 0.617 | 6.410 | 0.250 |
| **MGMT promoter methylation** | UM v. M | 0.970 | 0.627 | 1.500 | 0.891 |
| **Corticosteroid use** | Yes v. No | 1.271 | 0.861 | 1.877 | 0.228 |
| **Methylation subtype** | G-CIMP low v G-CIMP high | 4.072 | 2.231 | 7.430 | 0.000 |
| | Codel v G-CIMP high | 0.417 | 0.127 | 1.373 | 0.150 |
| | PA-like v G-CIMP high | 3.609 | 0.789 | 16.514 | 0.098 |

Supplementary table 3

| Gene.name | log2FoldChange |
|---|---|
| AJ011932.1 | -2.737589267 |
| PCDHGB4 | -2.571945205 |
| CEACAM5 | -2.5341973 |
| SHOX2 | -2.492427446 |
| KRT13 | -2.460839239 |
| AC104051.2 | -2.403250008 |
| IGHV1-18 | -2.330831346 |
| HOXA5 | -2.323982392 |
| LINC01956 | -2.300130233 |
| LINC02513 | -2.24479318 |
| NTS | -2.194539237 |
| SIX6 | -2.175101096 |
| EN1 | -2.158765071 |
| HOXA7 | -2.157135213 |
| MAGEA6 | -2.071106151 |
| HOXA10 | -2.059261084 |
| IL21-AS1 | -2.042870083 |
| IBSP | -2.036361159 |
| SCGB3A2 | -2.030888599 |
| CCDC198 | -2.018895459 |
| MTCYBP18 | -1.988541182 |
| HOXD9 | -1.97664222 |
| H1-9P | -1.974360534 |
| LINC01993 | -1.969448233 |
| AC015909.3 | -1.905007912 |
| KRT16 | -1.891557269 |
| HOXA9 | -1.883361185 |
| TBX5 | -1.867429903 |
| H19 | -1.801177989 |
| LRRC18 | -1.777074293 |
| DLK1 | -1.727130873 |
| IDO1 | -1.72137244 |
| CNIH3-AS1 | -1.719623402 |
| AC096669.1 | -1.707462577 |
| SCNN1B | -1.680229048 |
| AC068308.1 | -1.675592245 |
| PLEKHS1 | -1.656709761 |
| DAO | -1.643293356 |
| MFAP2 | -1.616778551 |
| HAGLROS | -1.612619442 |
| LINC01235 | -1.611484946 |
| HOXA4 | -1.603222266 |

| | |
|---|---|
| TBX5-AS1 | -1.597762594 |
| CHI3L1 | -1.591438123 |
| LHX5 | -1.575761551 |
| MIR4527HG | -1.52646164 |
| LINC01485 | -1.515218265 |
| AC062021.1 | -1.501396095 |
| TRIM71 | -1.490465977 |
| SLC17A8 | -1.479718763 |
| AC253536.4 | -1.460186616 |
| LINC01571 | -1.454158719 |
| LCE1E | -1.450018917 |
| NOX3 | -1.44740241 |
| AC104574.2 | -1.432576878 |
| IL13RA2 | -1.426872513 |
| C3orf22 | -1.426816188 |
| C2orf91 | -1.424551675 |
| AL390755.2 | -1.417799432 |
| LINC01579 | -1.410814502 |
| PCAT4 | -1.408801892 |
| IGF2-AS | -1.407277847 |
| IGF2BP3 | -1.394810745 |
| AC008080.4 | -1.388244005 |
| ASB11 | -1.374324083 |
| AL158058.1 | -1.37079861 |
| AC084864.1 | -1.360024733 |
| SPOCD1 | -1.356355284 |
| HAMP | -1.353158198 |
| TGFB2 | -1.350160759 |
| TRPC7 | -1.349144827 |
| APOH | -1.34872969 |
| DLGAP1-AS5 | -1.345754514 |
| SHISAL2B | -1.340049125 |
| ALPK2 | -1.338269773 |
| AC005999.1 | -1.335162176 |
| SRD5A2 | -1.332787731 |
| AC009097.2 | -1.332505081 |
| AC007402.1 | -1.327565511 |
| COL9A1 | -1.327235013 |
| AEBP1 | -1.309900005 |
| SLC34A2 | -1.302526748 |
| AC006372.3 | -1.29796104 |
| BARHL1 | -1.296094493 |
| PLAC8 | -1.287968261 |
| LINC01387 | -1.271777633 |
| AC061992.2 | -1.271585912 |
| AL354811.1 | -1.262808675 |
| NKX2-1 | -1.258751209 |

| | |
|---|---|
| AC008760.2 | -1.257935042 |
| ASB5 | -1.256795604 |
| PCDHGA3 | -1.252469346 |
| FGFBP2 | -1.250388105 |
| CXCL10 | -1.250319136 |
| SLPI | -1.239113683 |
| LINC00606 | -1.230027579 |
| IL21R | -1.229097284 |
| TFAP2A | -1.227993409 |
| IFI6 | -1.218666285 |
| S100A3 | -1.217530644 |
| ARHGAP36 | -1.214660604 |
| AL049839.2 | -1.214006748 |
| NA | -1.209976594 |
| PIRT | -1.206189984 |
| VGF | -1.202116275 |
| AC126773.4 | -1.200656398 |
| TRPM8 | -1.198024209 |
| SLCO4A1-AS1 | -1.192055585 |
| SLC18A1 | -1.191695755 |
| RPS3AP5 | -1.18708919 |
| AC073389.2 | -1.183287408 |
| AP005202.1 | -1.182697026 |
| TREM1 | -1.176000574 |
| ITK | -1.172545553 |
| ULBP1 | -1.166387637 |
| HNRNPKP3 | -1.162965555 |
| AC107419.1 | -1.162344351 |
| ARSF | -1.150883317 |
| LINC02574 | -1.150231164 |
| LCE1D | -1.143358674 |
| AC112493.1 | -1.14299355 |
| ABCA13 | -1.131583881 |
| PTX3 | -1.124334869 |
| C10orf105 | -1.121615634 |
| LINC02282 | -1.120792665 |
| GRHL3-AS1 | -1.119249705 |
| AC002546.1 | -1.119126233 |
| TIMP4 | -1.107649912 |
| FAM151A | -1.097568155 |
| NR1H4 | -1.096213209 |
| AC004899.2 | -1.091653154 |
| MYBL2 | -1.087380276 |
| INSM1 | -1.086795726 |
| SLC14A2 | -1.080179565 |
| CA9 | -1.078361897 |
| SPATA3 | -1.077455875 |

| | |
|---|---|
| ADAMTS7P4 | -1.075366136 |
| AC027281.2 | -1.071532654 |
| ADIG | -1.070317119 |
| LINC01224 | -1.070240332 |
| LINC02029 | -1.069804734 |
| NA | -1.067618019 |
| TMSB15A | -1.067419295 |
| GNLY | -1.063935711 |
| PCDHA7 | -1.059616824 |
| CFAP77 | -1.048266211 |
| CDC14C | -1.040485693 |
| LINC01349 | -1.03912138 |
| AC023421.1 | -1.034292301 |
| TNFSF13B | -1.029345357 |
| THEM7P | -1.027307556 |
| MT1F | -1.02610821 |
| LINC02777 | -1.021269904 |
| VEPH1 | -1.018627595 |
| EYA1 | -1.018248945 |
| USP30-AS1 | -1.011199972 |
| C21orf62 | -1.003904261 |
| MYO15A | 1.002369696 |
| REM1 | 1.036824656 |
| SLCO4C1 | 1.068178932 |
| AC103681.2 | 1.075356234 |
| GRM2 | 1.080043134 |
| CALML3-AS1 | 1.085922855 |
| OTOF | 1.090196171 |
| SGCG | 1.092779196 |
| TNIP3 | 1.104033991 |
| PRKCG | 1.129143448 |
| GNG13 | 1.135341651 |
| FBXO40 | 1.163078138 |
| TPO | 1.188794917 |
| DLX1 | 1.190730282 |
| CLEC4G | 1.194118952 |
| PCDHGB3 | 1.251510045 |
| S100A7 | 1.294257769 |
| MTCO3P12 | 1.341898853 |
| SLC22A9 | 1.369452854 |
| PRND | 1.413607686 |
| PRTN3 | 1.419613663 |
| KLK7 | 1.424761311 |
| AC140125.2 | 1.497638207 |
| CLEC4GP1 | 1.562749558 |
| SLC38A4 | 1.603508363 |
| NPIPB13 | 2.072398191 |

Supplementary table 4

| Gene.name | log2FoldChange |
|---|---|
| MTRNR2L1 | -5.520724037 |
| LINC01055 | -4.380363929 |
| PITX1 | -3.992410842 |
| CCDC198 | -3.91584956 |
| TFAP2B | -3.824382791 |
| DAO | -3.704166761 |
| AC068308.1 | -3.243847134 |
| CNN2P8 | -3.148696973 |
| AC023421.1 | -2.916224137 |
| LINC01485 | -2.808094538 |
| SLC14A2 | -2.798813886 |
| STON1-GTF2A1L | -2.725619928 |
| CCL1 | -2.706869245 |
| EN1 | -2.62447316 |
| ISL2 | -2.597031297 |
| CHIT1 | -2.530243103 |
| IL13RA2 | -2.505016251 |
| ENOX1-AS2 | -2.447615266 |
| AC022498.1 | -2.390691554 |
| CHRM5 | -2.390294807 |
| PCDHGB1 | -2.328605085 |
| PCDHGB6 | -2.312015563 |
| SLC14A1 | -2.270870574 |
| NMUR2 | -2.270733359 |
| SLC47A2 | -2.262412923 |
| AL603840.1 | -2.248180985 |
| MTCYBP18 | -2.224655459 |
| KIF6 | -2.223589491 |
| TFCP2L1 | -2.193455866 |
| LINC02308 | -2.167770915 |
| AL033519.1 | -2.136495258 |
| AC091151.1 | -1.975602204 |
| AC005999.1 | -1.965555734 |
| LINC01894 | -1.912781222 |
| LINC01579 | -1.9039876 |
| AL355916.2 | -1.84043909 |
| SFRP2 | -1.839939629 |
| GSX2 | -1.835719986 |
| CCL4L2 | -1.794607717 |
| NA | -1.778943819 |
| CRLF1 | -1.773109087 |
| LINC01235 | -1.758861682 |
| AC026316.3 | -1.736707567 |

| | |
|---|---|
| AL161935.1 | -1.732663246 |
| TNFSF13B | -1.723565612 |
| RN7SKP23 | -1.713341854 |
| AL355482.1 | -1.708920378 |
| TIGIT | -1.697341592 |
| HIRAP1 | -1.695118366 |
| LINC00994 | -1.693595145 |
| CCL4 | -1.689420884 |
| CCL3L1 | -1.685923314 |
| DLGAP1-AS5 | -1.665873392 |
| SNORC | -1.6651562 |
| IL1B | -1.661839939 |
| THORLNC | -1.652046554 |
| CCL3 | -1.647463936 |
| AC092040.2 | -1.643428448 |
| TNFSF18 | -1.635271455 |
| AC139491.2 | -1.630242068 |
| AC068790.1 | -1.594647512 |
| PGM5P4 | -1.588087948 |
| SALL4 | -1.576421413 |
| LPL | -1.574161491 |
| USH1C | -1.56772355 |
| AC092112.1 | -1.565593909 |
| FAM151A | -1.564868904 |
| AC091435.2 | -1.556627129 |
| FAM181A-AS1 | -1.546233218 |
| AC004485.1 | -1.533405325 |
| Z84468.1 | -1.522085163 |
| CCT7P2 | -1.521992916 |
| REELD1 | -1.507029939 |
| AP003472.1 | -1.501948058 |
| AP000424.1 | -1.486411807 |
| AL355974.2 | -1.477589802 |
| CALN1 | -1.471320426 |
| GFAP | -1.469268554 |
| LINC01736 | -1.443494141 |
| TRDN | -1.437372297 |
| AC005162.2 | -1.400400857 |
| TLR4 | -1.399146112 |
| SLC11A1 | -1.389398943 |
| AP004782.1 | -1.386916969 |
| AL390755.1 | -1.382538868 |
| AL391845.2 | -1.373598494 |
| BTC | -1.371751893 |
| AC084880.1 | -1.366963829 |
| AC084880.3 | -1.363921455 |
| LINC01117 | -1.361051862 |

| | |
|---|---|
| ACKR4 | -1.349531264 |
| FAM184B | -1.343255695 |
| ACOT11 | -1.34158337 |
| CH25H | -1.340510059 |
| MIR3151 | -1.331716185 |
| LINC01132 | -1.312126547 |
| AC093305.1 | -1.302303524 |
| AL035665.1 | -1.29310018 |
| LINC01933 | -1.260115378 |
| LINC01480 | -1.257478371 |
| SLC35E1P1 | -1.250371519 |
| TEKT3 | -1.216363248 |
| TMEM72 | -1.210581484 |
| ADGRE4P | -1.209793588 |
| CFAP300 | -1.200305634 |
| HOGA1 | -1.198967863 |
| GREB1L | -1.188824146 |
| LINC01094 | -1.181797216 |
| S100Z | -1.177196503 |
| WARS2-IT1 | -1.176083234 |
| PCBP3-AS1 | -1.175294791 |
| ELN-AS1 | -1.169291966 |
| LINC01354 | -1.154376228 |
| TPRG1-AS1 | -1.151270415 |
| LINC00877 | -1.139144855 |
| L3MBTL4-AS1 | -1.12221505 |
| AC093627.7 | -1.120734298 |
| LINC02145 | -1.106081339 |
| AL157823.2 | -1.098547658 |
| RHBDL3 | -1.09053745 |
| GYG2 | -1.075122004 |
| LRRC37A7P | -1.070713931 |
| KCNJ16 | -1.024858397 |
| CAVIN3 | 1.009076484 |
| LINC02761 | 1.018693732 |
| CSAG1 | 1.059398456 |
| ADAMTSL5 | 1.069855548 |
| HPGD | 1.12645928 |
| FGF18 | 1.143058008 |
| PLEKHG4 | 1.149908695 |
| MYL9 | 1.1897534 |
| TFPI | 1.2233101 |
| FOXS1 | 1.236144832 |
| COX4I2 | 1.249191043 |
| SPON2 | 1.28821644 |
| PXDNL | 1.404933681 |
| FSCN2 | 1.444151007 |

| | |
|---|---|
| AL109615.3 | 1.446914523 |
| FMO1 | 1.492060237 |
| CNN1 | 1.558405106 |
| GJA5 | 1.635759646 |
| SV2C | 1.67050452 |
| MYOCD | 1.781451983 |
| DES | 1.845422778 |
| LRRC36 | 1.969650897 |
| MYH11 | 2.157843483 |
| PCDHGA6 | 2.7791885 |
| SLC22A8 | 2.917933903 |
